# Stem cells actively suppress regenerative plasticity in human colon

**DOI:** 10.1101/2025.07.13.664589

**Authors:** Joris H. Hageman, Defne Yalçin, Julian R. Buissant des Amorie, Sascha R. Brunner, Thomas A. Kluiver, Aleksandra Balwierz, Franziska L. Langner, Maria C. Puschhof, Yannik Bollen, Thanasis Margaritis, Hugo J.G. Snippert

## Abstract

Insights into intestinal stem cell functioning during homeostasis and repair have been predominantly derived from genetic mouse models. This is in stark contrast to the largely unexplored situation in the human gut, where the underlying mechanisms and stimuli that induce regeneration are poorly understood. Here, we developed genetic strategies to characterize fluorescently labelled *LGR5*^+^ stem cells in normal human colon organoids. In parallel, we made diphtheria toxin-mediated cell type ablation compatible with human cells, thereby enabling in-depth interrogation of the sequence of events during depletion and reappearance of stem cells. Following *LGR5*^+^ stem cell depletion, most of the remaining epithelial cells entered a regenerative state characterized by fetal-like expression programs. Strikingly, this regenerative response was already initiated before stem cell loss, indicative of active communication between functional stem cells and progeny during homeostasis. We identified inactivation of retinoid X receptor (RXR) as a crucial trigger to initiate the regenerative response in colonocytes, with human colon stem cells being the producer of the RXR stimulus retinoic acid. Thus, stem cell-derived retinoic acid actively suppresses the regenerative state in colonocytes, explaining how surviving cells sense stem cell loss.

## Introduction

Intestinal stem cells (ISCs) provide a continuous supply of newly differentiated cells in a tissue with rapid cellular turnover^1^. Upon damage, which can be caused by inflammation, infection or cancer therapy, the ISC compartment can quickly recover to restore the integrity of the intestinal epithelial layer^2,3^. Genetic mouse models have been instrumental to uncovering extensive plasticity in response to acute ISC damage, where virtually every intestinal cell type shows capacity to revert to the stem cell state to restore homeostasis^4–6^. During this process, cells first adopt a transient regenerative state characterized by expression of a fetal-like gene expression program, before completing the process of dedifferentiation into adult ISCs^7–10^. However, the signals triggering cellular plasticity in response to ISC loss remain to be discovered and it is unclear whether all cell types have equal potential to respond. It has also yet to be established whether similar phenomena occur in the human colon, where cancer therapies broadly affect the stem cells. To date, it has been challenging to study stem cell regeneration in normal human organoids, due to several reasons such as paucity of effective genetic models.

The ISC marker *LGR5* is one of the most well-known marker genes for adult stem cells. Pioneering studies using genetic mouse models for visualization, lineage tracing and depletion of stem cells were based on *Lgr5* and have revolutionized our insights into stem cell functionality *in vivo*^1,2^. Human organoid models closely recapitulate cellular composition and physiology of *in vivo* epithelia, with human colon serving as the pioneering example^11^. Despite the increased application of CRISPR-mediated genome editing, the number of successful attempts to visualize and manipulate stem cells by complex genetic knock-ins in normal human organoids remains limited^12^, hindering studies into cellular functions and regenerative plasticity with high temporal resolution. Here, we generated human *LGR5* knock-in models akin to genetic mouse models, enabling combined visualization and depletion of human colon stem cells. We use these models to uncover the regenerative cell fate transitions in the normal human colon during recovery from stem cell loss.

## Results

### Depletion of *LGR5*^+^ human colon stem cells

To explore genetic knock-in (KI) strategies that maximize reporter expression from the *LGR5* locus, we developed different KI designs for targeted insertion at the C-terminus of *LGR5* (Figure S1A). In an *LGR5*-expressing colon cancer cell line (LS174T), the most robust expression levels were observed using a KI design with a bicistronic message separating mNeonGreen (Neon) from LGR5 by means of an internal ribosomal entry site (IRES), combined with a WPRE that stimulates protein yield (Figure S1B-D). Next, we copied the optimal KI strategy to normal human colon organoids, which we isolated and cultured in niche-inspired culture conditions^13^ to ensure budding organoid morphologies with crypt-like structures reminiscent of the compartmentalized colonic epithelium *in vivo* (Figure S2A,B). To trigger the targeted depletion of *LGR5*^+^ cells from the organoids, the genetic KI design included an inducible caspase 9 (iCasp9)^14,15^ (Figure S3A). Correct integration of the knock-in was confirmed (Figure S3B) and nuclear Neon fluorescence was largely confined to the expected location in organoid crypts (Figure S3C).

Unexpectedly, when iCasp9 was activated using an excess amount of activating dimerizer (AP), no cell killing was observed (Figure 1A, Figure S3D). In contrast, our KI approach was functional upon integration in a different locus (*KRT20*, Figure 1A, Figure S3E), as well as in LS174T cells with high *LGR5* expression (Figure S3F-H). To confirm that inducing apoptosis requires high expression levels of iCasp9, we boosted *LGR5* expression in our KI organoids with CHIR and Valproic Acid (VPA)^16^, which enabled us to achieve cell death induction in the stem cell population with the highest Neon-expression (Figure 1B, Figure S3I). These results show that under standard culture conditions, iCasp9-mediated cell type depletion is not functional when driven from the endogenous *LGR5* locus in normal human colon organoids, likely due to its low expression levels.

**Figure 1.**
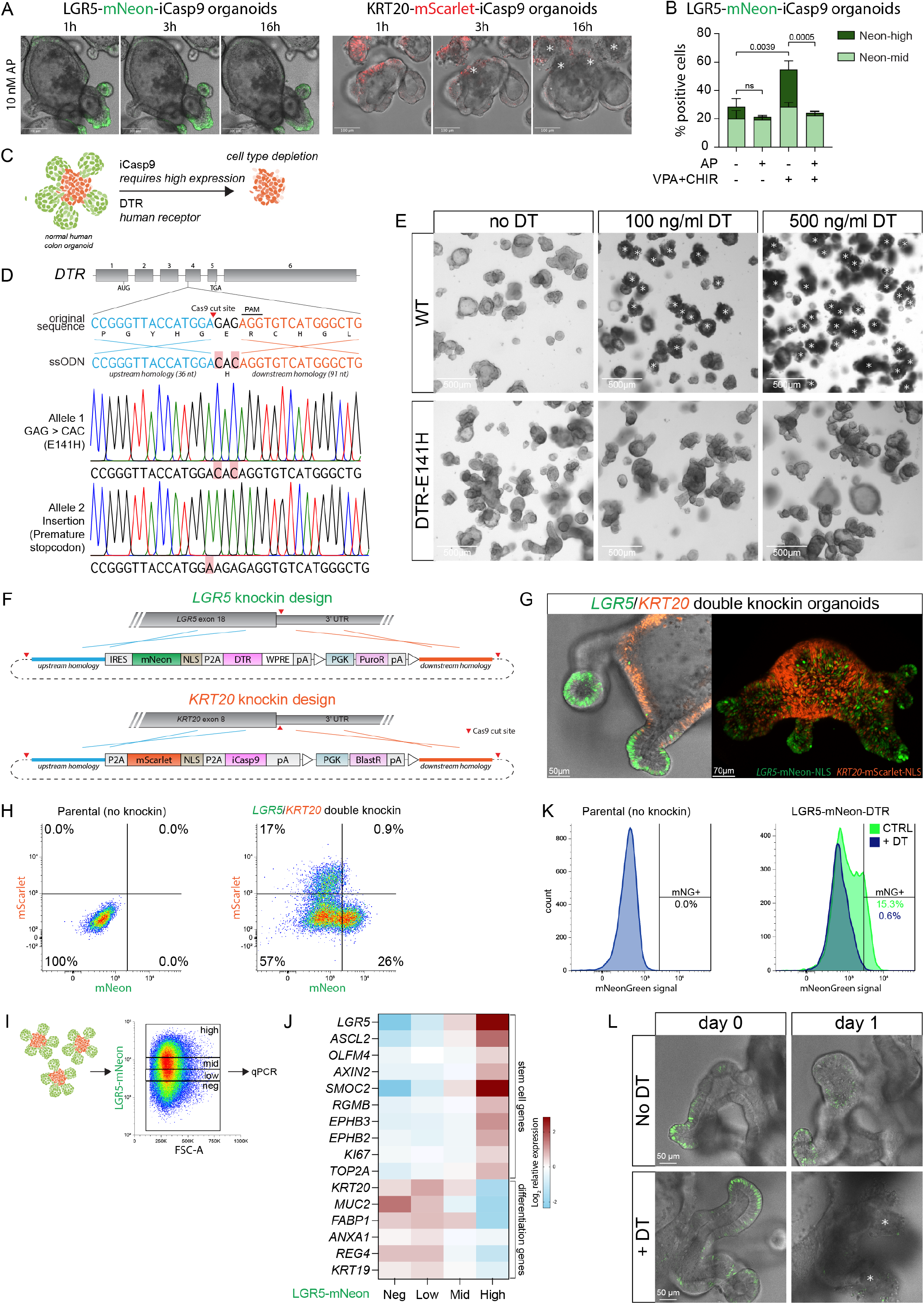
Visualization and depletion of *LGR5*^+^ and *KRT20*^+^ cells in normal human colon organoids. **(A)** Live cell microscopy of LGR5-mNeon-iCasp9 (left) and KRT20-mScarletI-iCasp9 (right) normal human colon organoids treated with 10 nM of the iCasp9 dimerizer AP20187 (AP). Asterisks indicate regions with dying cells. mNeon: mNeonGreen, iCasp9: inducible Caspase 9. Representative examples from *n* = 3 independent experiments. **(B)** Flow cytometry of the percentage of mNeon^+^ cells following different treatments of normal human colon organoids. AP only kills cells when the *LGR5* expression is first boosted with Valproic Acid (VPA) and CHIR, but even then, only the cells with the highest *LGR5* expression are successfully depleted. Values are corrected for the negative population to correct for the dying cells. Data are mean ± SD. ANOVA on the mNG-high population with Tukey’s correction, p-values are indicated. **(C)** Schematic representation of stem cell depletion (green cells). iCasp9-mediated cell type depletion requires high expression and is not functional at the *LGR5* locus. DTR-mediated cell type depletion provides an alternative, but DTR (HBEGF) is naturally expressed in all human colonic cells and therefore must first be inactivated. DTR: diphtheria toxin receptor. **(D)** CRISPR/Cas9 strategy to inactivate the naturally present DTR by introducing an E141H mutation. Organoids were co-electroporated with Cas9 targeting the *HBEGF* gene (*DTR*) and an ssODN with the desired mutation and homology regions. Sanger sequencing confirmed the presence of E141H on one allele and a knockout on the other allele. ssODN: single-stranded oligodeoxynucleotide **(E)** Brightfield imaging of wildtype and DTR-E141H normal human colon organoids following treatment with Diphtheria Toxin (DT). Asterisks indicate dying organoids. Representative examples of *n* = 3 independent experiments. **(F)** CRISPR/Cas9 knockin designs to generate LGR5/KRT20 double-knockin normal human colon organoids using in-trans paired Cas9 targeting. The upstream and downstream homology arms are flanked by guide sequences to enhance the knockin efficiency. *LGR5*^+^ and *KRT20*^+^ cells can be visualized and targeted independently due to different fluorescent proteins (mNeon, mScarletI) and depletion systems (DTR, iCasp9). IRES: internal ribosome entry site, mNeon: mNeonGreen, NLS: nuclear localization signal, WPRE: WHP Posttranscriptional Response Element, pA: poly A sequence, PGK: PGK promoter, PuroR: Puromycin Resistance cassette, BlastR: Blasticidin Resistance cassette. **(G)** Fluorescent microscopy stills of LGR5/KRT20 double knockin normal human colon organoids. Left: maximum projection of 3 Z planes of fluorescent channels and 1 Z plane of brightfield channel. Right: Full organoid maximum Z projection of fluorescent channels. **(H)** Flow cytometry of LGR5/KRT20 double knockin normal human colon organoids. Representative examples of *n* = 3 independent experiments. **(I)** FACS strategy to sort cells based on LGR5-mNeonGreen expression levels for qPCR. **(J)** Gene expression of cells with different expression levels of LGR5-mNeon, sorted from normal human colon organoids as shown in h. Expression of stem cell markers and differentiation markers was analyzed with qPCR. **(K)** Flow cytometry of Diphtheria Toxin (DT) treatment of normal human colon organoids with LGR5-mNeon-DTR. Representative example of *n* = 3 independent experiments. **(L)** Live cell microscopy stills of LGR5-mNeon-DTR normal human colon organoids following DT treatment. Asterisks indicate dying *LGR5*^+^ buds.

Therefore, we turned to an alternative cell type depletion strategy that is widely used in genetic mouse models for its high sensitivity, which relies on the ectopic expression of Diphtheria Toxin Receptor (DTR)^2,17^. Unfortunately, this strategy is not compatible with human organoid models, because DTR is in fact the ubiquitously expressed human membrane-anchored glycoprotein HB-EGF that acts as the natural entry factor for Diphtheria Toxin (DT)^18^ (Figure 1C). To desensitize human colon organoids to DT, we mutated HB-EGF in its DT-binding domain by one amino acid substitution (E141H), phenocopying the non-sensitive mouse variant^19,20^ (Figure 1D). As intended, while wildtype organoids completely died upon DT administration, organoids containing “murinized” HB-EGF^E141H^ were insensitive even to high concentrations of DT (Figure 1E).

In these HB-EGF^E141H^ organoids, we generated double knock-ins with LGR5-Neon-DTR and KRT20-mScarlet-iCasp9 to simultaneously visualize *LGR5*^+^ stem cells and *KRT20*^+^ differentiated cells and enable their independent cell type depletion (Figure 1F). Correct integrations and expression patterns were confirmed (Figure S4A-C), with *LGR5*^+^ cells localizing to the bottom of crypt structures, while *KRT20*^+^ cells were restricted to the body of the organoids (Figure 1G,H, Video S1). Moreover, LGR5-Neon^+^ cells co-expressed other known ISC markers (Figure 1I,J), while KRT20-mScarlet^+^ cells expressed markers of differentiation (Figure S4D). Importantly, low concentrations of DT were sufficient to successfully kill the entire LGR5-Neon^+^ cell population within a day (Figure 1K,L, Video S2). To our knowledge, this is the first time that DTR-mediated cell type depletion is successfully applied in a human setting.

### Rapid reappearance of stem cells

To study the regenerative response of human colonic epithelium to acute stem cell loss, we administered a 6-hour pulse of DT to the LGR5/KRT20 KI organoids, which triggered the depletion of all *LGR5*^+^ cells within the next 20 hours (Figure 2A). Using live-cell microscopy and flow cytometry, we observed a remarkably rapid onset of robust cellular plasticity and reappearance of *de novo LGR5*^+^ cells within 24 hours after stem cell depletion (48 hours after first DT exposure) (Figure 2B,C, Video S3). Following passaging (72 hours post DT), fully restored organoids formed that appeared identical to the initial condition, with the expected *LGR5*^+^/*KRT20*^+^ cell type ratios (Figure 2D,E). Reappearance of *LGR5*^+^ cells was not an artefact of DT treatment, as *de novo LGR5*-Neon expression was not observed in an KI organoid line without DTR in the *LGR5* locus, nor upon sublethal DT concentrations in DTR-containing organoids (Figure S5A,B). In contrast, depletion of the differentiated cell compartment using KRT20-iCasp9 was not directly followed by reappearance of *KRT20*^+^ cells, but rather by expansion of the remaining *LGR5*^+^ stem cell compartment (Figure 2F-G, Video S4). Upon passaging, these organoids also restored to their initial state (Figure 2H). To confirm that *de novo LGR5*^+^ cells emerged from regenerative plasticity of *LGR5*^-^ cells, rather than from multiplication of rare residual *LGR5*^+^ cells, we performed cell tracking analysis of live-cell imaging recordings (Figure 2I, Video S5,S6). The resulting tracking traces validated that reappearing *LGR5*^+^ cells originated from *LGR5*^-^ cells (Figure 2J). Considering that stem cell identity *in vivo* depends on niche-derived factors, we tested the influence of various signaling pathways relevant to the ISC compartment. Withdrawal of signaling ligands following stem cell depletion confirmed that stem cell reappearance in human colonic organoid epithelia is highly dependent on external stimulation by niche factors, most notably through WNT activation and BMP suppression (Figure 2K).

**Figure 2.**
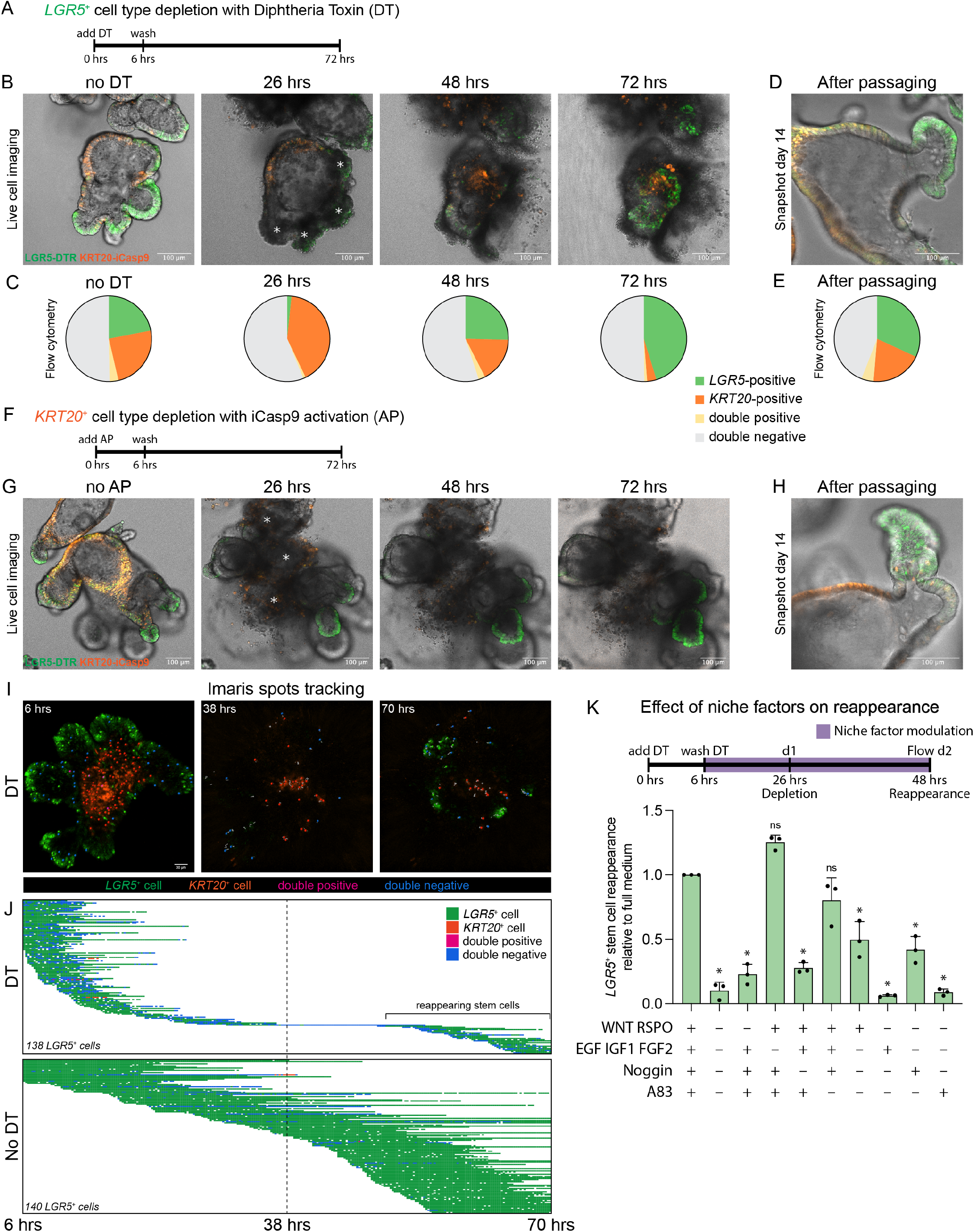
*LGR5*^+^ cells reappear following their ablation in normal human colon organoids. **(A)** Experimental design to deplete *LGR5*^+^ cells with DT and examine their reappearance. Normal human colon organoids were treated with a 6-hour pulse of DT to deplete *LGR5*^+^ cells and allow them to reappear. **(B)** Live cell microscopy stills showing *LGR5*^+^ cell killing and reappearance. Asterisks indicate dying *LGR5*^+^ buds. Representative images of *n* = 3 independent experiments. **(C)** Quantification of *LGR5*^+^ cell killing and reappearance with flow cytometry. Representative example of *n* = 6 independent experiments. **(D)** Representative microscopy still of an organoid that was passaged 3 days after DTR-mediated stem cell depletion and subsequently grown for 11 days.EXT **(E)** As in D, quantified with flow cytometry. **(F)** Experimental design to deplete *KRT20*^+^ cells with inducible caspase 9 (iCasp9), which is activated by AP20187 (AP). Normal human colon organoids were treated with a 6-hour pulse of AP to deplete *KRT20*^+^ cells. **(G)** Live microscopy stills showing *KRT20*^+^ cell killing and subsequent *LGR5*^+^ cell compartment expansion. Representative images of *n* = 3 independent experiments. **(H)** Representative microscopy still of an organoid that was passaged 3 days after AP-mediated *KRT20*^+^ cell type depletion and subsequently grown for 11 days. **(I)** Tracking of individual *LGR5*^+^ cells (green dots) during stem cell depletion and reappearance. **(J)** Tracking traces of individual *LGR5*^+^ cells during live cell microscopy of LGR5/KRT20 knockin normal human colon organoids with or without a 6-hour DT pulse. Each horizontal line represents a single cell. Colors of the traces indicate *LGR5* and *KRT20* status based on fluorescence of mNeon and mScarlet, respectively. **(K)** Flow cytometry quantification of *LGR5*^+^ cell reappearance (2 days after DT pulse) in the indicated medium conditions, displayed relative to the level of reappearance in full medium. Data are mean ± SD. *n* = 3 independent experiments. ANOVA with Dunnett’s correction comparing each condition to full medium, *p<0.001.

### Stem cell loss triggers regenerative state

To gain temporal insight into the cellular states and signaling pathways that are activated during the regenerative process upon stem cell depletion, we performed single-cell RNA sequencing (scRNA-seq) at successive timepoints following the DT pulse (Figure 3A,B). Unsupervised clustering revealed the expected cell types of the human colonic epithelium with their corresponding marker genes, including stem cells, goblet cells, colonocytes, enteroendocrine cells and tuft cells (Figure 3C,D). Reassuringly, stem cells were not detected 24 hours following the DT pulse, confirming that LGR5-

**Figure 3.**
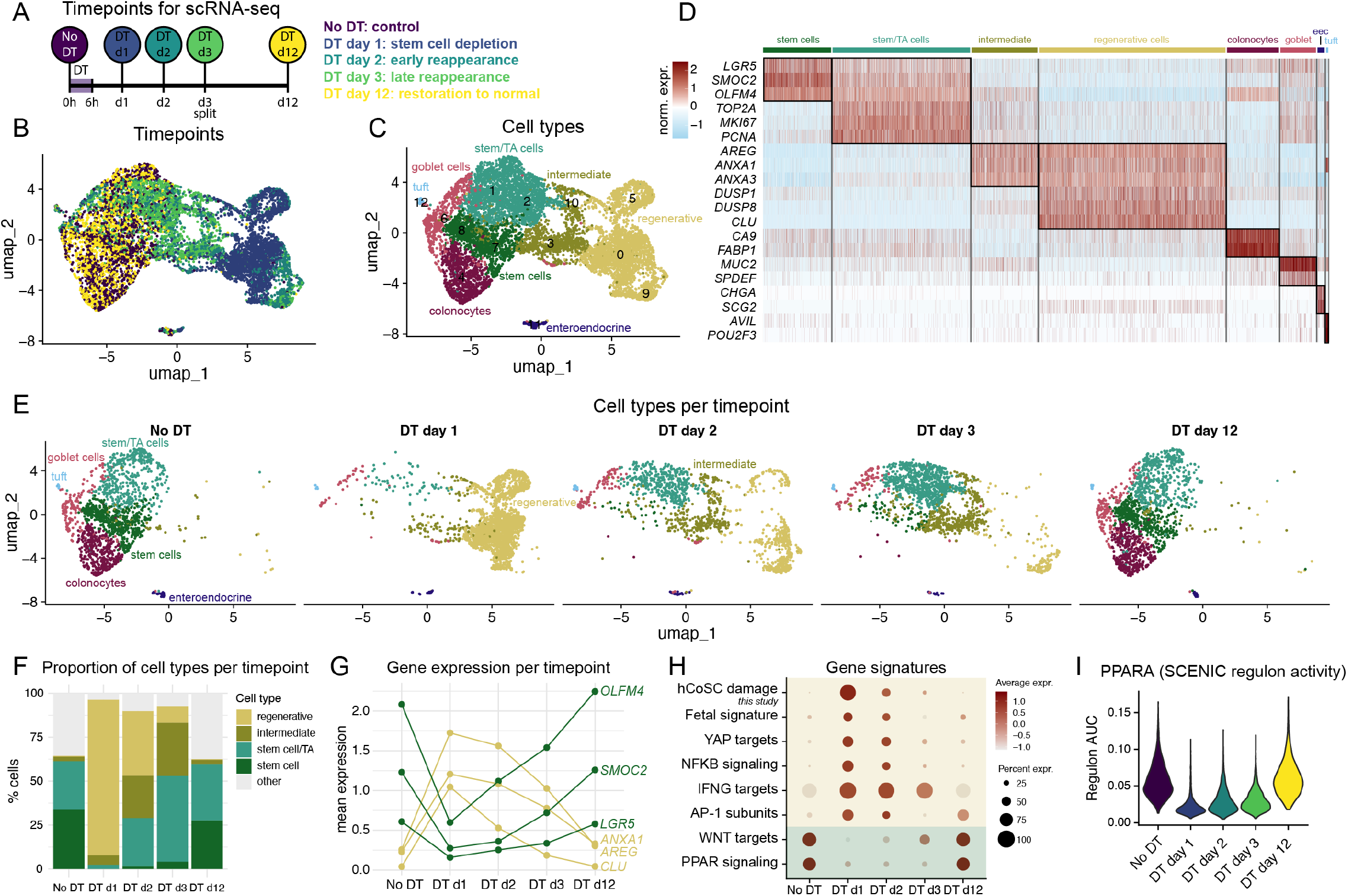
Stem cell depletion induces a regenerative state in the differentiated cells. **(A)** Experimental set-up for single cell RNA sequencing (scRNA-seq) during stem cell depletion and reappearance in normal human colon organoids. Organoids were harvested for scRNA-seq at the indicated timepoints before or after a pulse of DT. **(B)** Dimensionality reduction (UMAP) of scRNA-seq dataset of normal human colon organoids before and after stem cell depletion, where colors indicate the timepoints as shown in panel A. **(C)** Same as B, where colors indicate the corresponding cell types. **(D)** Heatmap showing the expression of marker genes of the indicated cell types per cell. Norm. expr: log2 normalized expression (row-scaled). **(E)** UMAP split by timepoint showing the abundancies of different cell types over time. **(F)** Relative percentage of cell types per indicated timepoints. **(G)** Pseudo-bulk gene expression of stem cell markers (green) and regenerative cell markers (yellow) at the indicated timepoints. **(H)** Dot plot showing the expression of the indicated gene signatures, displayed as average expression over all cell types per timepoint. Signatures were derived from refs^8,51–54^, KEGG, or this study (human colon stem cell [hCoSC] damage) (see Table S2). **(I)** Violin plot of PPARA regulon activity (AUC) over time during stem cell depletion and reappearance, as calculated by SCENIC.

Neon-DTR is a faithful marker of the stem cell compartment and effectuates its full depletion when triggered (Figure 3E,F). Interestingly, the majority of the remaining cells had transitioned to a regenerative state almost simultaneously with cell death execution in the stem cells, characterized by expression of YAP target genes (such as *AREG* and *CLU*) and genes associated with a fetal-like expression program (Figure 3D-F, Figure S6A). During day 2 and day 3 after the initial insult, cells gradually adopted an intermediate phenotype between the regenerative and homeostatic state and, concomitantly, the absolute number of reappearing stem cells increased (Figure 3E-G). Following passaging and outgrowth of regenerated cultures (DT d12), organoids fully restored and they were largely indistinguishable from the initial condition (Figure 3B,E,F). Next, we derived a gene signature that is activated upon acute damage to stem cells in human colon (hCoSC damage signature) through differential expression analysis between the samples ‘DT day 1’ (stem cell depletion) and ‘No DT’ (Table S1). In addition to the hCoSC damage signature, regenerative cells showed upregulation of NFKB signaling, IFNG targets, AP1 subunit expression, and downregulation of canonical WNT and PPAR signaling (Figure 3H, Figure S6B). Colonocytes displayed the highest activity of the PPAR signaling pathway during homeostasis (Figure S6C). SCENIC analysis^21^, which integrates co-expression patterns with cis-regulatory motifs to infer transcription factor (TF) activity, confirmed the increased activity of AP-1, as well as the reduced activity of PPARA in the regenerative state (Figure 3I, Figure S6D,E). We noticed that surviving cells completely reversed their phenotype to the regenerative state almost immediately after perturbation of stem cell function (DT kills by blocking all protein synthesis). This strikingly fast response suggests a previously unrecognized form of proactive communication during homeostasis, in which healthy stem cells actively prevent dedifferentiation in surrounding differentiated cells.

### TFs associated with the regenerative state

To identify the TFs and epigenetic states involved in the major cell state shifts upon acute stem cell loss, we performed combined profiling of single-cell transcriptome and chromatin state (10x multiome) at 24 hours and 36 hours following DT pulse and on the No DT control. We confirmed the presence of all expected cell types with their corresponding markers (Figure 4A,B) and we detected the emergence of the same regenerative gene programs as in the scRNA-seq dataset, including our hCoSC damage signature, immediately following stem cell loss (Figure 4C). We then analyzed the scATAC-seq module to identify TF motif enrichment across the cell types (Figure 4D). Regenerative cells showed significant accessibility enrichment to AP-1 members (such as JUN and FOSB) as well as NFKB (REL and RELA) motifs (Figure 4D, Figure S6F). In contrast, chromatin regions associated to the retinoic X receptor (RXR) and its partners such as PPARA, which during homeostasis shows the highest activity in colonocytes, became inaccessible following DT (Figure 4E,F).

**Figure 4.**
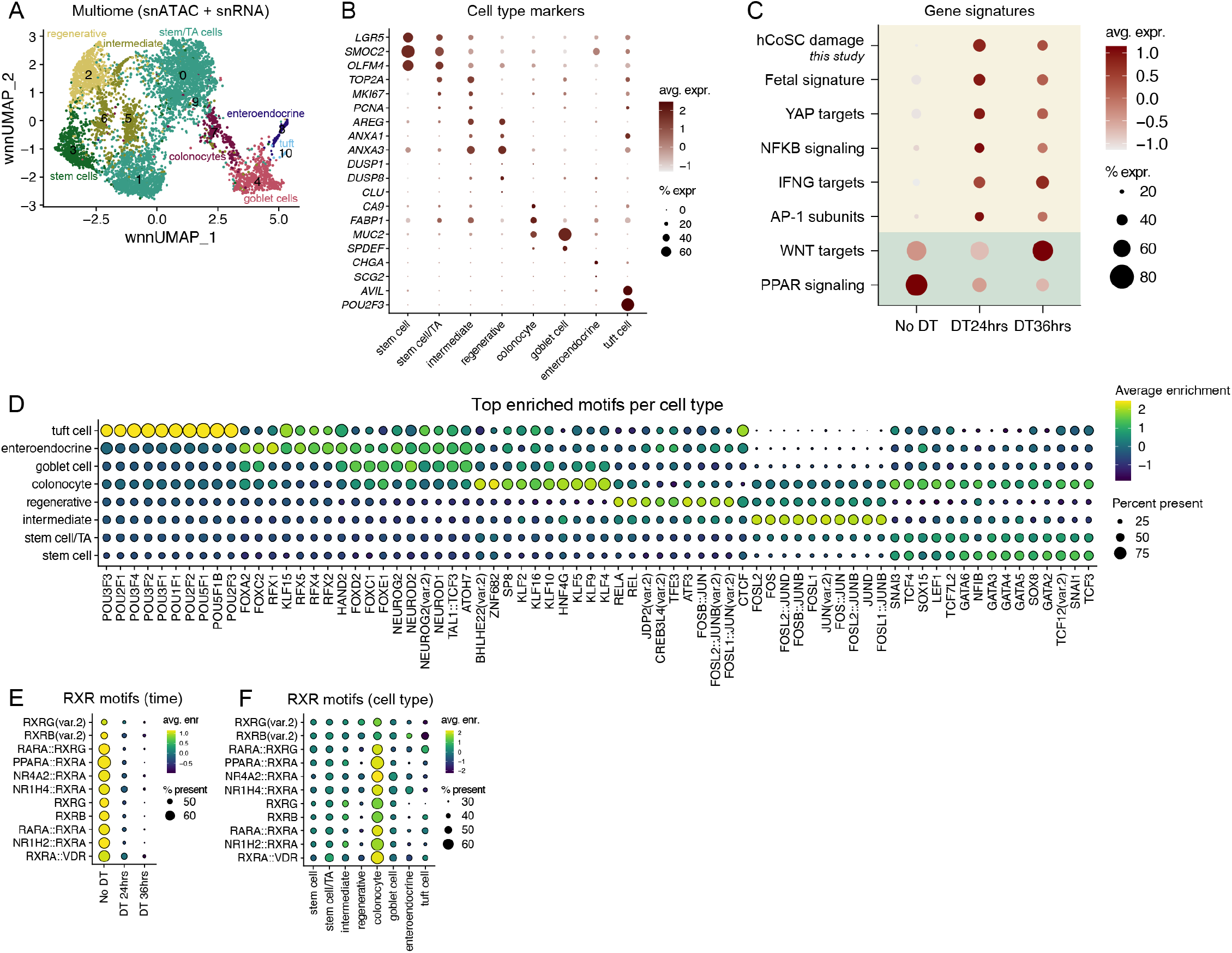
Combined scRNA-seq and scATAC-seq analysis reveals transcription factors linked to the regenerative state. **(A)** Weighted nearest neighbor (wnn) UMAP with unsupervised clustering of single nucleus (sn)ATAC-seq and snRNA-seq (10x multiome) dataset colored by cell type. Analyzed samples are No DT, 24 hrs after DT, and 36hrs after DT. **(B)** Dot plot showing normalized RNA expression of indicated marker genes per cell type in the multiome dataset. **(C)** Dot plot showing the expression of the indicated gene signatures (the same signatures as in Figure 3H), displayed as average expression over all cell types per timepoint in the multiome dataset. Green: signatures relating to homeostasis, yellow: signatures relating to the regenerative state. **(D)** Dot plot showing the top enriched motifs per cell type based on ATAC-seq (ChromVar). **(E)** Dot plot showing the enrichment of RXR-related motifs over time. **(F)** Dot plot showing the enrichment of RXR-related motifs per cell type.

### RA from stem cells suppresses regeneration

The near-immediate fate reversion in almost all colonocytes upon perturbation of stem cell function suggests stem cells communicate a ‘keep-calm-and-carry-on’ signal to ensure homeostatic conditions. We reasoned that retinoic acid (RA) could be a candidate for that signal, because colonic stem cells specifically express ALDH1^22–24^, which is the rate-limiting enzyme that converts retinol into RA. As shown above, the activity of the RA receptor RXR is highest in the colonocytes of unperturbed organoids, which is in line with earlier reports that RXR activity promotes colonocyte differentiation^25,26^. Suppression of dedifferentiation by RA could therefore explain why plasticity in colonocytes is immediately triggered once stem cell function is compromised.

To test this hypothesis, we generated organoids with a CRISPR-mediated mScarlet knock-in at the regeneration marker *ANXA1*^8,27^ (Figure 3G, Figure 5A, Figure S7A). Although low baseline levels of *ANXA1* can be detected in most cell types under normal conditions, regenerative cells are distinctly marked by high expression of *ANXA1* (Figure 5B, Figure S7B,C). Following manipulation of an array of regeneration-related signaling pathways in normal human colon organoids, we observed a large increase in regenerative *ANXA1*^high^ cells following treatment with the RXR antagonist HX531 (referred to as RXRi) (Figure 5C,D, Figure S7D). RXRi also induced upregulation of several other genes associated with the regenerative state (Figure 5E). Similarly, inhibition of the RA-catalyzing enzyme ALDH1 increased the number of *ANXA1*^high^ cells (Figure 5F), demonstrating that epithelial production of RA is involved in the suppression of the regenerative state. To identify the source of RA production, we examined our scRNA-seq dataset and performed fluorescent immunostaining for ALDH1A1, both of which revealed its enrichment in stem cells (Figure 5G, Figure S7E). We then used AldeFluor, a fluorescent substrate of ALDH enzymes, to document the spatial pattern of RA production within organoids. Consistent with a previous report showing that colonic stem cells from normal human gut can be isolated with AldeFluor^22^, we noticed strong enrichment of AldeFluor at the bottom of crypt-like structures in normal human colon organoids, confirming that RA production is confined to the stem cell niche (Figure 5H). To confirm that RA signals a ‘keep-calm-and-carry-on’ message to colonocytes, we administered excess RA at the moment of stem cell depletion and observed that it was able to counteract regenerative plasticity (Figure 5I). Conversely, we had noticed that exit from the regenerative state is in turn accompanied by activation of RXR and canonical WNT signaling (Figure 3H,I). Accordingly, stem cell reappearance was suppressed by the continuous presence of RXR inhibition, locking the cells in the regenerative state (Figure 5I). Taken together, our data support a model in which stem cell damage in human colon organoids leads to inactivation of RXR signaling due to the loss of stem cell-derived RA, thereby triggering regenerative reprogramming in surrounding colonocytes (Figure 6).

**Figure 5.**
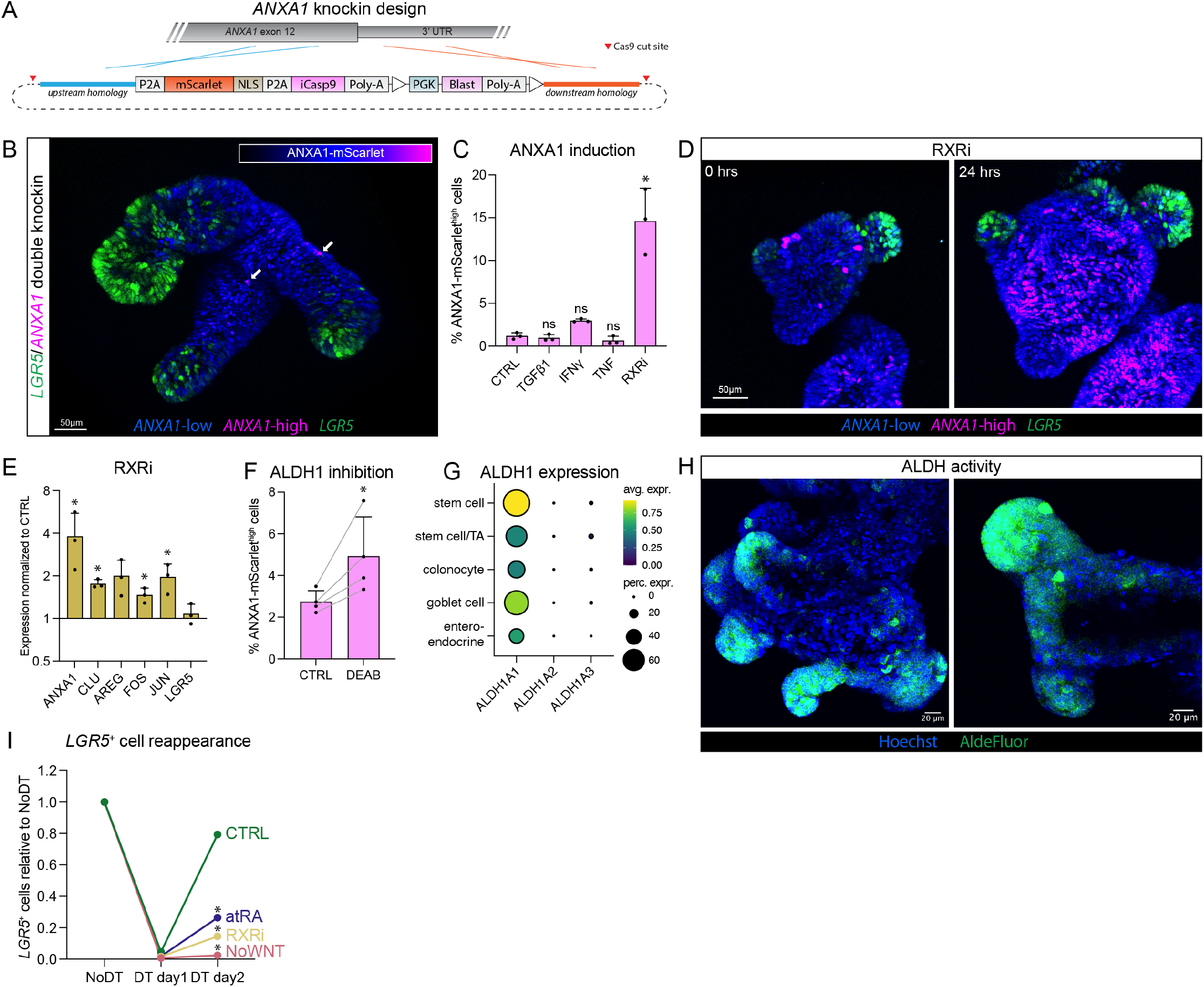
Stem cell-derived retinoic acid suppresses the regenerative state. **(A)** CRISPR knockin design for ANXA1-mScarlet-iCasp9 to generate LGR5/ANXA1 double knockin normal human colon organoids. **(B)** Representative microscopy still (maximum Z projection) of normal human colon organoid with LGR5/ANXA1 double knockin showing a small number of cells with high ANXA1 expression (magenta, indicated with arrows). **(C)** Flow cytometry of ANXA1-mScarlet knockin normal human colon organoids to identify factors that induce the presence of *ANXA1*^high^ cells (See Figure S7C for FACS gating). RXRi: RXR inhibition with the RXR antagonist HX531. Data are mean ± SD. *n* = 3 independent experiments. ANOVA with Dunnett’s correction comparing each condition to the control, *p<0.0001. **(D)** Microscopy stills of LGR5/ANXA1 double knockin organoids before or 24 hrs after addition of RXRi (HX531). Representative example of *n* = 3 independent experiments. **(E)** Gene expression following 24 hrs of RXRi (HX531) treatment in normal human colon organoids, normalized to control without RXRi (qPCR). Data are mean ± SD. *n* = 3 independent experiments. Ratio paired t test, *p<0.05. **(F)** Flow cytometry of ANXA1-mScarlet organoids 24 hrs after treatment with the ALDH1 inhibitor DEAB. Data are mean ± SD. *n* = 4 independent experiments. Paired t test, *p=0.048. **(G)** Dot plot showing the expression of the different ALDH1 genes in our scRNA-seq dataset of normal human colon, per cell type. **(H)** Microscopy images (maximum Z projection) of the AldeFluor assay (green) in normal human colon organoids. Two representative organoids are shown. **(I)** Flow cytometry of *LGR5*^+^ cell reappearance in different conditions following a 6-hour DT pulse. The proportion of *LGR5*^+^ cells is displayed relative to the number of *LGR5*^+^ cells before DT. atRA: all-trans Retinoic Acid, RXRi: HX531, NoWNT: withdrawal of WNT. *n* = 3-6 independent experiments. ANOVA with Dunnett’s correction comparing all conditions on day 2 to the CTRL on day 2, *p<0.05.

**Figure 6.**
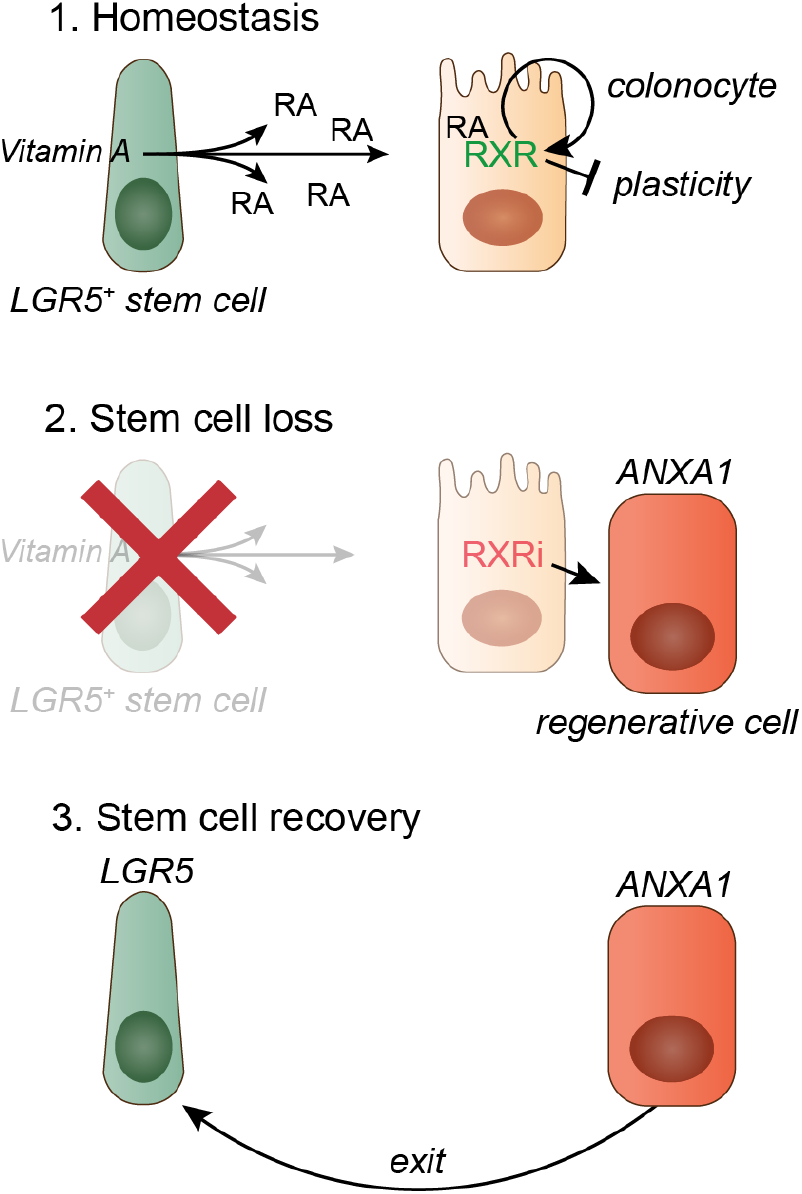
Model for the induction of regenerative plasticity in the human colon. During homeostasis, intestinal stem cells secrete RA, preventing regenerative plasticity in colonocytes. Upon stem cell loss, RA production ceases and colonocytes activate the regenerative cell state. These regenerative cells subsequently give rise to *de novo LGR5*^+^ stem cells. RA: retinoic acid, RXR: retinoid X receptor.

## Discussion

Although there is an extensive body of research into intestinal stem cell functionality in health and disease using mouse models, insights into human stem cell dynamics remain limited. This is partially because widely used marker genes, such as the intestinal stem cell marker LGR5, are generally lowly expressed, hindering their utility as drivers of molecular tools like iCasp9. To address this, we present a novel strategy to visualize and selectively deplete cell types in human model systems, based on the favored DTR-mediated cell type depletion method that thus far was limited to mouse models. The complex multi-purpose gene knock-ins that we applied to human colon models, are applicable to virtually all cell types with well-defined marker genes, as well as pluripotent stem cell-derived tissues such as cerebral organoids, greatly expediting functional studies into unique human cell types. While iCasp9 effectuates cell killing faster than DTR, DTR-mediated cell killing is less reliant on expression levels, rendering it more suitable for endogenous gene knock-in strategies to ablate specific cell type populations in their entirety.

We used our knock-in models to investigate the rapid recovery of the stem cell compartment upon DTR-mediated damage and how surviving cells in the epithelium sense stem cell loss. Unlike the prevailing model where the regenerative response is triggered by damage signals, we discovered that regenerative plasticity in human colon is actively suppressed during homeostasis by communication between stem cells and colonocytes (Figure 6). Specifically, stem cells continuously signal a ‘keep-calm-and-carry-on’ message to colonocytes by means of the RXR ligand RA. Upon stem cell damage, the RA signal is lost, and regenerative plasticity is unleashed in the colonocytes. This regenerative state is characterized by previously described fetal-like expression programs^28^ and activation of YAP, AP-1, NF-κB, and IFN-γ signaling. We derived a specific hCoSC damage signature that can be utilized by future studies to identify acute stem cell damage responses in various *in vivo* settings. A related mechanism has been reported in the skin, where RA maintains hair follicle stem cell lineage commitment, and its transient loss following epidermal damage enables a switch in cell fate^29^.

We anticipate that multiple signals, in addition to diminished RA signaling, can act together to achieve the full regenerative response. For example, changes in mechanical properties of the surrounding matrix can activate regenerative signaling programs ^30^ and signaling factors that are induced in damage settings such as TGF-β^31^ have been shown to promote the regenerative state. While abrogated RXR activity represents an early trigger that communicates stem cell loss, it remains to be resolved how multiple signals converge to drive the complete regenerative cascade. Similarly, the exact details and order of molecular events that drive exit from the regenerative state warrants further investigation.

Intriguingly, in mouse small intestinal organoids, RXR inhibition has been shown to stabilize the regenerative state, but was unable to trigger it from a homeostatic state^26^. A key discrepancy is the different localization of RA biosynthesis. Unlike human colon, *Aldh1a1*, the rate-limiting enzyme of RA production, is expressed outside of the stem cell compartment in mouse intestinal enterocytes. It is unclear why mouse small intestine utilizes different triggers of regeneration than human colon, but it does highlight the importance of studying epithelial cell fate transitions in human tissue models.

A multitude of studies have investigated the cell of origin of newly generated stem cells in the murine intestine following stem cell loss^4,5,32^. Together, these studies established that virtually every cell type, including enterocytes and various secretory cells, have the capacity to dedifferentiate into stem cells. In addition, reserve stem cells and non-cycling stem cells have also been implicated in the regenerative response^33–35^. In the human intestine, quiescent stem cells^12^ and the extremely rare tuft cells^36^ have been identified as sources of regenerating cells. Although we cannot formally prove which cell type is most potent to replenish lost stem cells, our data strongly suggest a prominent role for the abundant colonocytes. First, colonocytes were the only cell type in the scRNA-seq dataset that completely converted following stem cell loss, and their high abundance corresponds to the high numbers of regenerative cells observed following stem cell loss. Second, colonocytes displayed the highest activity of RXR signaling during homeostasis, indicating that the regenerative trigger of RXR inhibition is primarily manifested in this population. Our new model systems provide a powerful platform to investigate the cellular dynamics underlying regeneration in human tissues and to elucidate its molecular mechanisms with high temporal resolution.

Of conceptual interest are the similarities between our findings and the plasticity-driven regenerative cell states in cancer, which are involved in therapy resistance and metastatic relapse^27,37,38^. Mechanistically, RXR activity has been reported to block the entry into a regenerative cancer state^39^, underscoring the similarities between normal tissue and cancer. Detailed insights into the molecular mechanism of cell fate plasticity can be instrumental to modulate regenerative processes in health and disease.

## Acknowledgements

We thank all members of the Snippert lab, in particular Sander Mertens and Suzanne van der Horst for their assistance with FACS sorting and Bas Ponsioen and Sander Mertens for their critical comments on the manuscript. We thank Joep Sprangers (UMC Utrecht) for advice on the CRISPR strategy. We thank Eric Kalkhoven and Suzanne van der Veen (UMC Utrecht) for reagents and helpful discussions. We thank Niels Groenen and Jeroen van Velzen from the FACS facility of the Princess Máxima center. We thank the Single-Cell Genomics facility of the Princess Máxima center for their assistance with single-cell transcriptomics and epigenomics analysis. We also thank the Cell Microscopy Core (CMC) of the Center for Molecular Medicine, UMC Utrecht for providing training and service for Imaris software. This work is part of the Oncode institute, which is partly financed by the Dutch Cancer Society. This work was supported by a Starting Grant from the European Research Council (ERC). This paper was typeset with the bioRxiv word template by @Chrelli: www.github.com/chrelli/bioRxiv-word-template

## Author contributions

Conceptualization: J.H.H., H.J.G.S., Investigation (cell culture, electroporation, FACS, microscopy, immunofluorescence, qPCR): J.H.H., D.Y., F.L.L., Generation of organoid models: J.H.H., J.R.B.d.A., CRISPR designs: J.H.H., M.C.P., Y.B., H.J.G.S., Single-cell transcriptomics and epigenomics: J.H.H., T.A.K., A.B., T.M., Analysis single-cell transcriptomics and epigenomics: J.H.H., J.R.B.d.A., S.R.B., T.A.K., Funding acquisition: H.J.G.S., Supervision: T.M., H.J.G.S., Writing: J.H.H., H.J.G.S.

## Competing interest statement

The authors declare no competing interests.

## Materials and Methods

### LS174T cell culture

LS174T cells (clone W4^40^) were cultured in RPMI-1640 (Sigma), supplemented with 10% FBS (Bodinco), 1% Penicillin/Streptomycin (P/S, Lonza), and 1% GlutaMAX (Gibco) at 37 °C under 5% CO_2_ atmosphere. Cells were passaged using Trypsin-EDTA (Sigma).

### Organoid culture

Normal human colon organoids were previously derived from the sigmoid colon of a 64-year-old male donor, and cultured in growth factor-reduced Matrigel (Corning) at 37°C (5% CO_2_ atmosphere), in niche-inspired culture medium that induces intestinal cell type diversity^13^, consisting of advanced DMEM/F12 (Gibco) supplemented with 1% P/S, 1% GlutaMAX, 1% HEPES buffer (Gibco), 10% Noggin conditioned medium (in-house production), 20% R-spondin1-conditioned medium (in-house production), 1x B27 (Gibco), 1.25 mM N-acetylcysteine (Sigma), 500 nM A83-01 (Tocris), 50 ng/ ml recombinant human EGF (PeproTech), 100 ng/ml recombinant human IGF-1 (BioLegend), 50 ng/ml recombinant human FGF-basic (FGF-2, Peprotech), and 0.5 nM Wnt surrogate (U-protein Express). Organoids were passaged with Trypsin-EDTA for 1-2 minutes at 37 °C. After organoids were trypsinized into single cells and small fragments, cells were embedded in 10 μL droplets of Matrigel which were solidified for 10 minutes at 37 °C, before adding culture medium which was supplemented with 10 μM ROCK inhibitor Y-27632 (Gentaur) for the initial 3-4 days after passaging.

For experiments, organoids were split to single cells and small fragments and plated at a high density (splitting ratio 1:8). Subsequently, organoids were replated (without breaking them apart) after 4 days at a ∼1:30 ratio (∼30 organoids/10 μL Matrigel) to allow them to expand further without spatial restriction by neighboring organoids. Experiments were performed 7-9 days after splitting (3-5 days after replating), when organoids presented as budding structures. During experiments, organoids were treated with 5 ng/ml DT (Sigma D0564) or 10 nM AP20187 (Sigma SML2838), unless different concentrations are indicated. Concentrations used for additional experiments are 10 ng/ml TGF-β1 (Immunotools 11343160), 20 ng/ml IFN-γ (Immunotools 11343536), 100 ng/ml TNF (Abcam ab259410), 2 μM HX531 (RXRi, Sigma SML2170), 100 μM DEAB (Sigma D86256), 10 μM all-trans Retinoic Acid (atRA, Stem Cell Technologies 72264). For the niche factor screen on reappearance following DT, full medium was used during the initial 6-hour DT pulse, after which growth factors were withdrawn as shown in the figure.

### CRISPR/Cas9 constructs

The design of the targeting vectors was based on our previously published targeting vectors^41^, and the vectors were modified as indicated in the figures using infusion cloning (Takara Bio) and Golden Gate cloning. All sequences were confirmed with Sanger sequencing or whole plasmid sequencing. In addition to locus-specific targeting vectors for *LGR5, KRT20*, and *ANXA1*, we developed generalized constructs (without locus-specific homology arms) that can be easily adopted to any desired locus with one single Golden Gate assembly, as we previously described^41,42^. To generate DTR-E141H organoids, we developed an ssODN and guide targeting the *HBEGF* gene, according to previously reported guidelines^43^. Cas9 plasmids with guide sequences were generated with wildtype Cas9 (LGR5, HBEGF, based on addgene 48139) or nickase Cas9 (KRT20, ANXA1, based on addgene 48141) as previously described^44^. Guide sequences and ssODN used are listed in Table S3.

### Transfection of LS174T cells

LS174T cells were transfected using 4.5 μL TransT-LT1 (Mirus), with 500 ng Cas9 guide and 1 μg targeting vector, which was added to the cells 2-3 days after splitting. After 3 days, 10 μg/ml puromycin was added to select for knockin cells.

### Electroporation of organoids

Electroporation of organoids was based on a previously published protocol^45^. Briefly, organoids were expanded for ∼8 days after splitting. 1.25% DMSO was added 2 hours before organoid harvest. Organoids were trypsinized for 1 minute to single cells and small fragments and were then washed with Opti-MEM (Gibco) and resuspended in 11 μg targeting vector, 4μg Cas9/ guide, added to 100 μL with BTXpress (Fisher). Cells were electroporated in a NEPA21 electroporator following described settings^45^, but with 190 V poring pulse. Organoids were then incubated for 10 minutes at room temperature after addition of 400 μL Opti-MEM with 10 μM ROCK inhibitor, before plating them in Matrigel and adding culture medium with ROCK inhibitor. After 3-4 days, antibiotic selection was performed with 1 μg/ml Puromycin or 4 μg/ml Blasticidin, after which the antibiotics were included in the culture medium for several passages. In case of using the wildtype Cas9, monoclonal organoid lines were derived by hand-picking individual organoids. Besides the knockin alleles, the secondary (untargeted) alleles were analyzed by Sanger sequencing to exclude the presence of indel mutations. Primers for amplifying genomic DNA are listed in Table S3. In the case of the introduction of the DTR-E141H mutation, the same protocol was followed except with 5 μg ssODN instead of targeting vector, and 10 μg instead of 5 μg Cas9/guide. Correctly modified organoids were selected by adding 5 ng/ml DT 4 days after electroporation for a duration of 4 days, after which monoclonal DT-resistant organoids were hand-picked and analyzed with Sanger sequencing for their DTR/HBEGF mutation.

The availability of the organoid lines that have been generated in this study is restricted by the Utrecht Medical University (UMC) Utrecht ethical committee. To receive these organoid lines, a request with the appropriate forms has to be made through this committee, which will determine if the request corresponds with the informed consent of the patient.

### Microscopy

LS174T cells were seeded onto a fibronectin-coated glass bottom plate for imaging. Organoids were seeded in Matrigel 1-3 days before imaging on an 8 chamber Ibidi glass bottom plate. Fluorescent imaging was performed on a Leica SP8 WLL scanning confocal microscope with Leica Application Suite X (37 °C, 6.8% CO_2_), with a temporal resolution of 15-30 minutes and Z resolution of 5 μm.

For analysis, FIJI (v2.14.0) and Imaris (v10.2) were used. In Imaris, we summed up the LGR5 (Neon) and KRT20 (mScarlet) channels to track nuclei with spots detection using the autoregressive motion algorithm. Therefore, only cells that expressed at least low levels of LGR5 or KRT20 could be tracked. We used 2D classification to identify *LGR5*^+^, *KRT20*^+^, double positive, and double negative cells. Data were analyzed in Rstudio (v.2024.12.1) and displayed with ComplexHeatmap (v2.20.0).

Brightfield images of organoids were captured on an EVOS M5000.

### Antibody staining

Fluorescent staining with antibodies was performed as previously described^46^. Primary antibodies (Ki67 ab15580 1:1000, CHGA sc-1488 1:500, MUC2 sc-15334 1:500, AldoB ab137628 1:500, KRT20 M701901-2 1:100, or

ALDH1A1 MA5-29023 1:100) were incubated at 4°C overnight. Secondary antibodies (Alexa Fluor goat and donkey) were incubated overnight at 4°C. Organoids were imaged on a Leica SP8 WLL scanning confocal microscope in clearing agent (60% glycerol and 2.5 mol/L fructose).

### FACS

LS174T cells or organoids were harvested and trypsinized similarly as for passaging. Flow cytometry was conducted on a BD FACS Celesta Cell

Analyzer. DAPI-negative cells were analyzed as the live cell compartment. Gates were determined based on negative control samples (parental lines without knockins and without DAPI staining) Flow cytometry data was analyzed and visualized using BD FACSDiva software and the free online tool https://floreada.io. FACS sort for qPCR was performed on a SONY MA900 with a 100 μm nozzle.

### qPCR

RNA was extracted with the Nucleospin RNA isolation kit (Macherey-Nagel 740955). cDNA was generated from RNA using the iScript cDNA Synthesis Kit (Bio-Rad). For the qPCR, 10 ng cDNA was mixed with 0.5 μM forward and reverse primer each and 5 μL PowerTrack SYBR Green (Applied Biosystems) per well. qPCR was performed on the QuantStudio 5 Real-Time PCR System and results were analyzed using the ΔΔCt method using *ACTB* and *PBGD* as reference genes. Primer sequences are listed in Table S3.

### scRNA-seq

Organoids containing LGR5-Neon-DTR were passaged on 5 different days (as described above), such that they could all be treated with a 6-hour DT pulse (5ng/ml) after 9 days of outgrowth (5 days after replating), and could then be harvested on the same day. Flow cytometry of aliquoted material of all samples was performed (as described above) to confirm stem cell depletion and reappearance based on *LGR5* expression. On the day of harvest for scRNA-seq, organoids were harvested with 5 μM EDTA in advanced DMEM/F12 to dissolve the Matrigel for 20 minutes at 4°C. Organoids were split into single cells with Trypsin-EDTA and filtered with a 20 μm strainer. For multiplexing, the 5 samples were incubated with 5 different Cell Multiplexing Oligos (CMOs) according to protocol CG000391 (10x Genomics), with prolonged incubation time (15 minutes) at room temperature. Dead cells were removed according to protocol CG000093 (10x Genomics) using the MACS dead cell removal kit (Miltenyi Biotec). ∼10,000 cells per sample were then pooled and loaded to the 10x Genomics Chromium Controller for droplet-based scRNA-seq. Libraries were sequenced on the NovaSeq sequencer (used for GEX, ∼32K reads/cell) and NovaSeq XPlus sequencer (used for CMO-lipid-based de-multiplexing, ∼8K reads/cell), both from Illumina.

### scRNA-seq analysis

Read alignment to GRCh38 and CMO-based demultiplexing of the samples was performed using CellRanger (v.7.1.0). This way, ∼2000 cells could be reliably assigned to each sample. Analysis of scRNA-seq data was performed using Seurat^47^ (v.5.0.1). Cells with <10% mitochondrial genes, >1000 unique features (genes), and >2000 counts (total features) were selected for further analysis. Data were then normalized and scaled using the functions NormalizeData and ScaleData. Unsupervised clustering was performed with FindNeighbors (20 dimensions, k.param = 20) and FindClusters (resolution 0.6). To define the cell types, markers for cell clusters were determined using FindAllMarkers (Table S4). The hCoSC damage signature was derived using FindMarkers for the top 100 positive markers (ranked on adjusted p value) between the samples DT day 1 and No DT. The AddModuleScore function was used to calculate gene expression signature scores, which were derived from this study, the indicated publications, and mSigDB. To determine regulon activity we used pySCENIC^21^ (v.0.11.2). We then used FindMarkers (Seurat) for No DT and DT day 1 to calculate differentially active regulons based on the SCENIC AUC scores.

### 10× multiome

Organoids of two donors containing LGR5-Neon-DTR were treated for 6 hrs with a pulse of 5ng/ml DT, and harvested at the indicated timepoints following the start of the DT pulse. Organoids were cryopreserved as whole organoids in CellBanker (AmsBio) for a maximum duration of 2 days at −150 °C. Organoids were then simultaneously thawed and processed for 10x multiome according to protocol CG000375 (10x Genomics). Samples relating to the same timepoints of the two donors were pooled. Nuclei were isolated in NP40 lysis buffer (Sigma 74385) with glass douncers. The samples were filtered using a 70 μm strainer. Nuclei were stained with 7-AAD (Invitrogen) and isolated using a SONY SH800S FACS sorter (100 μm nozzle). Then, nuclei were permeabilized and their open chromatin tagged with transposase. Transposed nuclei were then loaded into Chromium X (10x Genomics). Single cell RNA and ATAC libraries were generated from the same cells according to protocol CG000338 Rev F (10x Genomics). Libraries were sequenced on the NovaSeq 6000.

### 10× multiome analysis

Raw data were processed using CellRanger ARC (v2.0.2). SNP-based demultiplexing was performed with Souporcell^48^ (v2.5) and analyzing male/ female-specific genes (*XIST, PRKY, LINC00278, USP9Y, UTY, TTTY14*). Reads were aligned using CellRanger (v.8.0.0) and analyzed using Seurat (v5.0.1) and Signac (v.1.14.0). Downstream analyses were performed on the samples derived from the same donor as in the rest of the manuscript. Cells were filtered as follows: 100 < nCount_ATAC < 40000 & 300 < nCount_ RNA < 8000 & percent.mt < 2 & TSS.enrichment > 3 & nucleosome_signal < 2. The RNA assay was normalized using NormalizeData and scaled using ScaleData. To define the cell types, markers for cell clusters were determined using FindAllMarkers (Table S5). Peak calling in the ATAC assay was performed using MACS2 (v2.2.9.1). Dimensionality reduction was performed with RunTFIDF, FindTopFeatures, and RunSVD with default settings. Multimodal analysis of both assays was performed with FindMultiModalNeighbors, RunUMAP, and FindClusters with resolution 0.8 and SLM algorithm. For motif analysis, ChromVar^49^ (v1.26.0) was used after adding the human JASPAR2020 motifs^50^. To determine the top enriched motifs per cell type or timepoint, we used FindAllMarkers on the ChromVar assay for positive values, using the logistic regression test, min. pct = 0.05 & latent.vars = ‘nCount_ATAC’. For the motifs for RXR and its partners, we analyzed all motifs that contained “RXR” in their motif name. The AddModuleScore function was used for the RNA assay to calculate gene expression signature scores with the RNA assay, similarly as for the scRNA-seq dataset. This included the hCoSC damage signature derived from the scRNA-seq dataset.

### AldeFluor assay

The AldeFluor assay (StemCell Technologies) was carried out according to the manufacturer’s protocol. Organoids cultured for 7 days were replated onto glass-bottom dishes and maintained for an additional 2 days. For the assay, organoids embedded in Matrigel were incubated at 37 °C for 35 minutes with the AldeFluor reagent and Hoechst 33342, in the presence or absence of the ALDH inhibitor DEAB. Organoids treated with DEAB served as a negative control to establish background fluorescence levels. Fluorescent imaging was performed using a Leica SP8 WLL scanning confocal microscope.

### Statistical analyses

Statistical analysis was performed as noted in the figure legends using GraphPad Prism (v10). Data distribution was assumed to be normal, but this was not formally tested. All statistical tests were two-tailed. P-values < 0.05 were deemed to be statistically significant.

### Data availability

No new code was developed for this manuscript. scRNA-sequencing data and 10x multiome data are available at Zenodo (https://doi.org/10.5281/zenodo.15405448). The availability of the cell lines that have been generated in this study is restricted by the Utrecht Medical University (UMC) Utrecht ethical committee. To receive these cell lines, a request with the appropriate forms has to be made through this committee, which will determine if the request corresponds with the informed consent of the patient.

## Supplementary information

**Video S1. LGR5-mNeon/KRT20-mScarlet double knock-in normal human colon organoids**.

Three representative examples of double knock-in organoids are shown. Green: LGR5-mNeon-NLS, orange: KRT20-mScarlet-NLS

**Video S2. Live cell microscopy of depletion of *LGR5***^**+**^ **cells from normal human colon organoids**.

DT was added at t=0 hrs inducing cell death in *LGR5*^+^ cells. Green: LGR5-mNeon-NLS.

**Video S3. Live cell microscopy of depletion and reappearance of *LGR5*^+^ cells in normal human colon organoids**.

At t=0 hrs, organoids were treated with a 6-hour pulse of DT to deplete *LGR5*^+^ cells and allow for their reappearance. Green: LGR5-mNeon-NLS, orange: KRT20-mScarlet-NLS.

**Video S4. Live cell microscopy of depletion of *KRT20***^**+**^ **cells in normal human colon organoids**.

At t=0 hrs, organoids were treated with a 6-hour pulse of AP (iCasp9 dimerizer) to deplete *KRT20*^+^ cells. Green: LGR5-mNeon-NLS, orange: KRT20-mScarlet-NLS.

**Video S5. Live tracking of depletion and reappearance of *LGR5***^**+**^ **cells in normal human colon organoids**.

At t=0 hrs, organoids were treated with a 6-hour pulse of DT to deplete *LGR5*^+^ cells and allow for their reappearance. Green nuclei: LGR5-mNeon-NLS, orange nuclei: KRT20-mScarlet-NLS. Green spots: *LGR5*^+^ cells, orange spots: *KRT20*^+^ cells, pink spots: double positive cells, blue spots: double negative cells.

**Video S6. Live tracking of normal human colon organoids**.

Control condition of Video S5 without DT. Green nuclei: LGR5-mNeon-NLS, orange nuclei: KRT20-mScarlet-NLS. Green spots: *LGR5*^+^ cells, orange spots: *KRT20*^+^ cells, pink spots: double positive cells, blue spots: double negative cells.

**Table S1. Differential expression analysis of DT day 1 vs No DT for hCoSC damage signature generation**.

DE analysis was performed on the scRNA-seq dataset between the stem cell depletion condition (DT day 1) and the control condition (No DT).

**Table S2. Gene signatures used in this study**.

**Table S3. Primers, guide sequences, and ssODN used in this study.**

**Table S4. Cell cluster markers in the scRNA-seq dataset**.

**Table S5. Cell cluster markers in the multiome dataset**.

**Figure S1.**
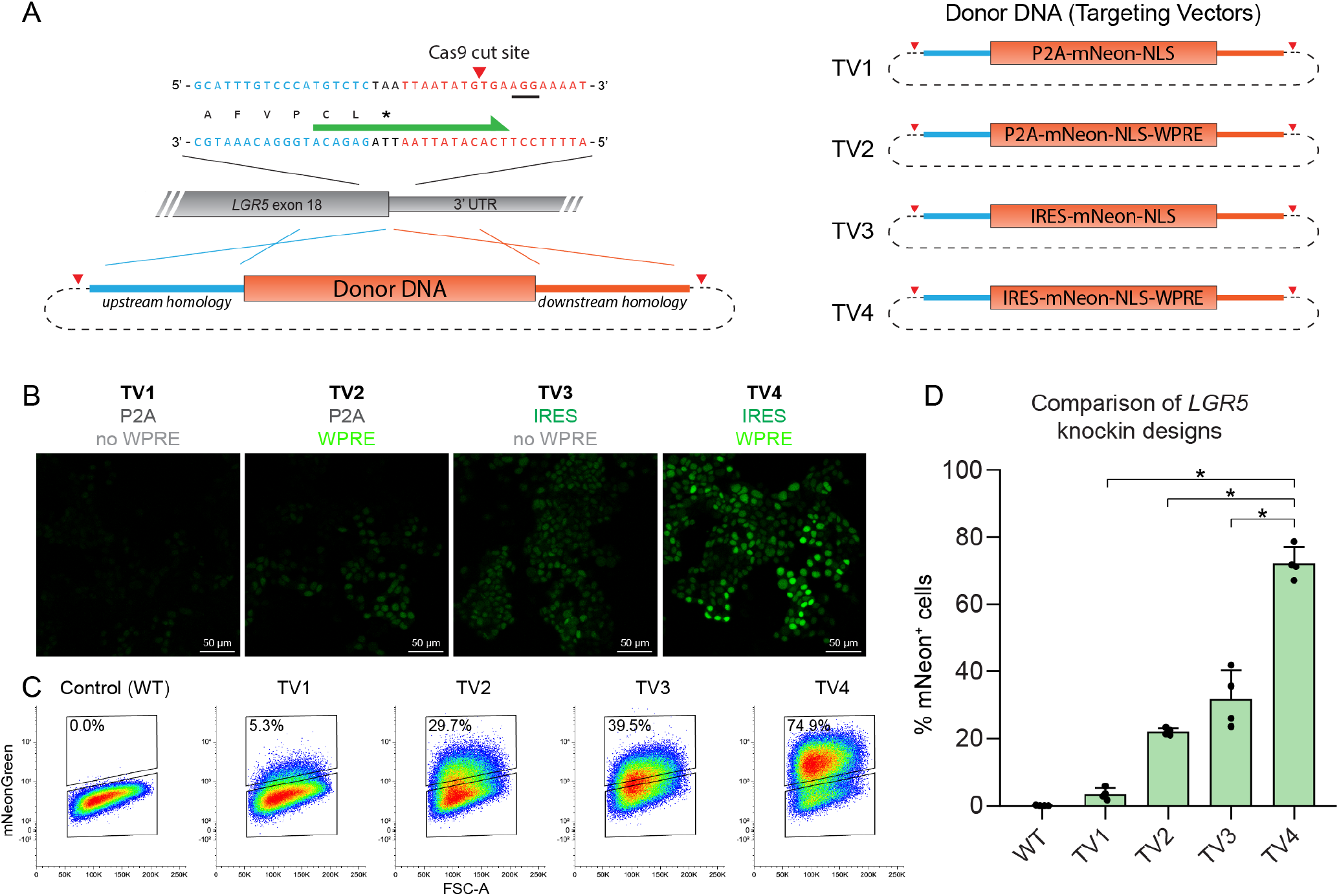
Optimization of *LGR5* knockin design. **(A)** CRISPR design to generate fluorescent knockins at the final exon of the *LGR5* locus using in-trans paired Cas9 targeting. The upstream and downstream homology arms are flanked by guide sequences to enhance the knockin efficiency. The fluorescent protein mNeonGreen is separated from the LGR5 protein using either a P2A (TV1, TV2) or IRES (TV3, TV4). A WPRE has been included in TV2 and TV4. TV: targeting vector, WPRE: WHP Posttranscriptional Response Element, IRES: internal ribosome entry site, NLS: nuclear localization signal. **(B)** Fluorescent microscopy to compare the different *LGR5* knockin approaches in LS174T cells. **(C)** Flow cytometry to compare the different *LGR5* knockin approaches in LS174T cells. **(D)** uantification of panel c. Data are mean ± SD. *n* = 4 independent replicates. ANOVA with Tukey’s correction. *p<0.0001.

**Figure S2.**
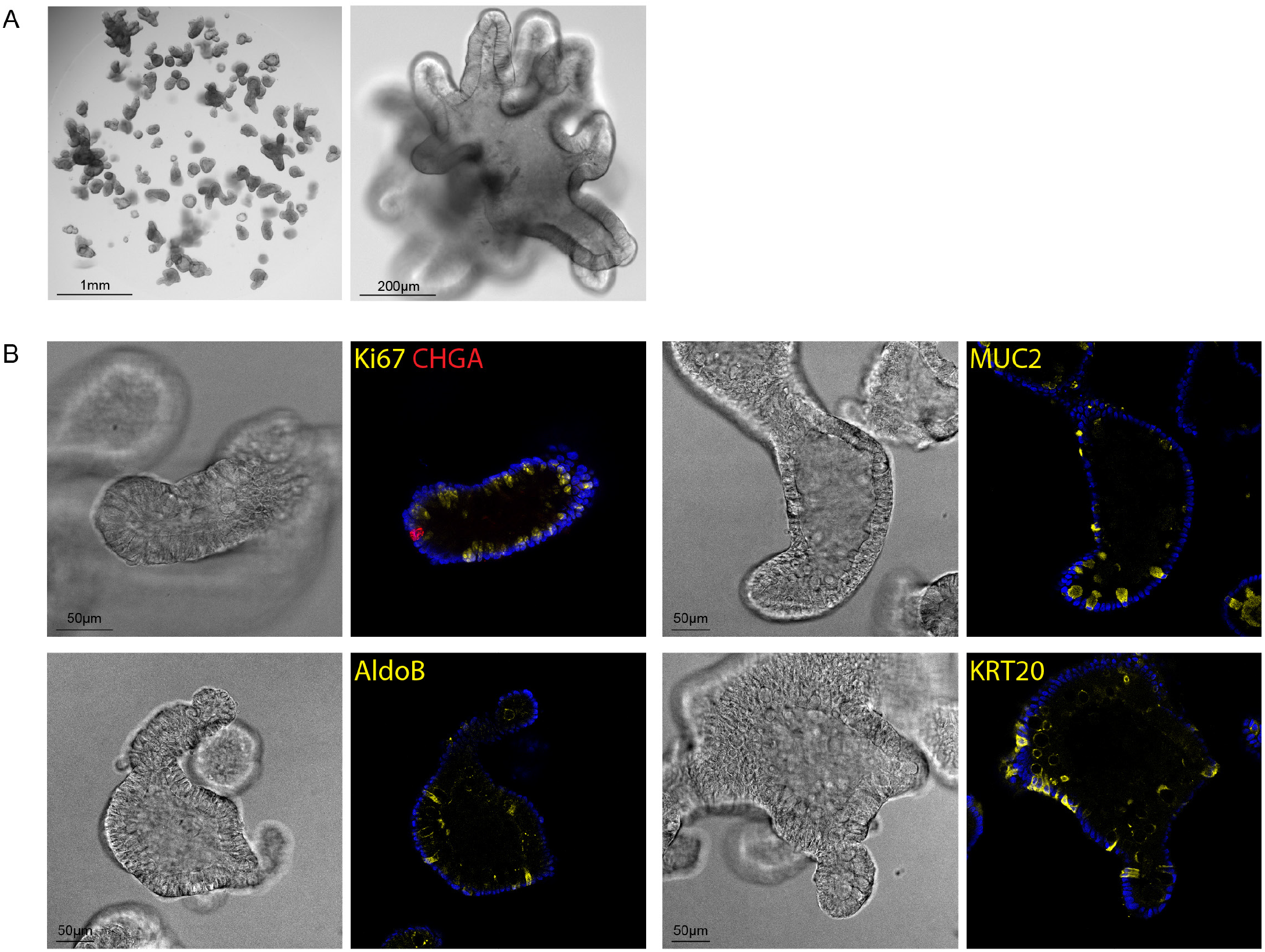
Normal human colon organoids grow as budding structures and contain a diversity of cell types. **(A)** Brightfield microscopy images of normal human colon organoids presenting with budding morphology. **(B)** Immunofluorescent staining for different markers of colonic epithelial cell types. Ki67: proliferating cells. Chromogranin A (CHGA): enteroendocrine cells. Mucin 2 (MUC2): goblet cells. Aldolase B (ALDOB): enterocytes. Keratin 20 (KRT20): differentiated cells.

**Figure S3.**
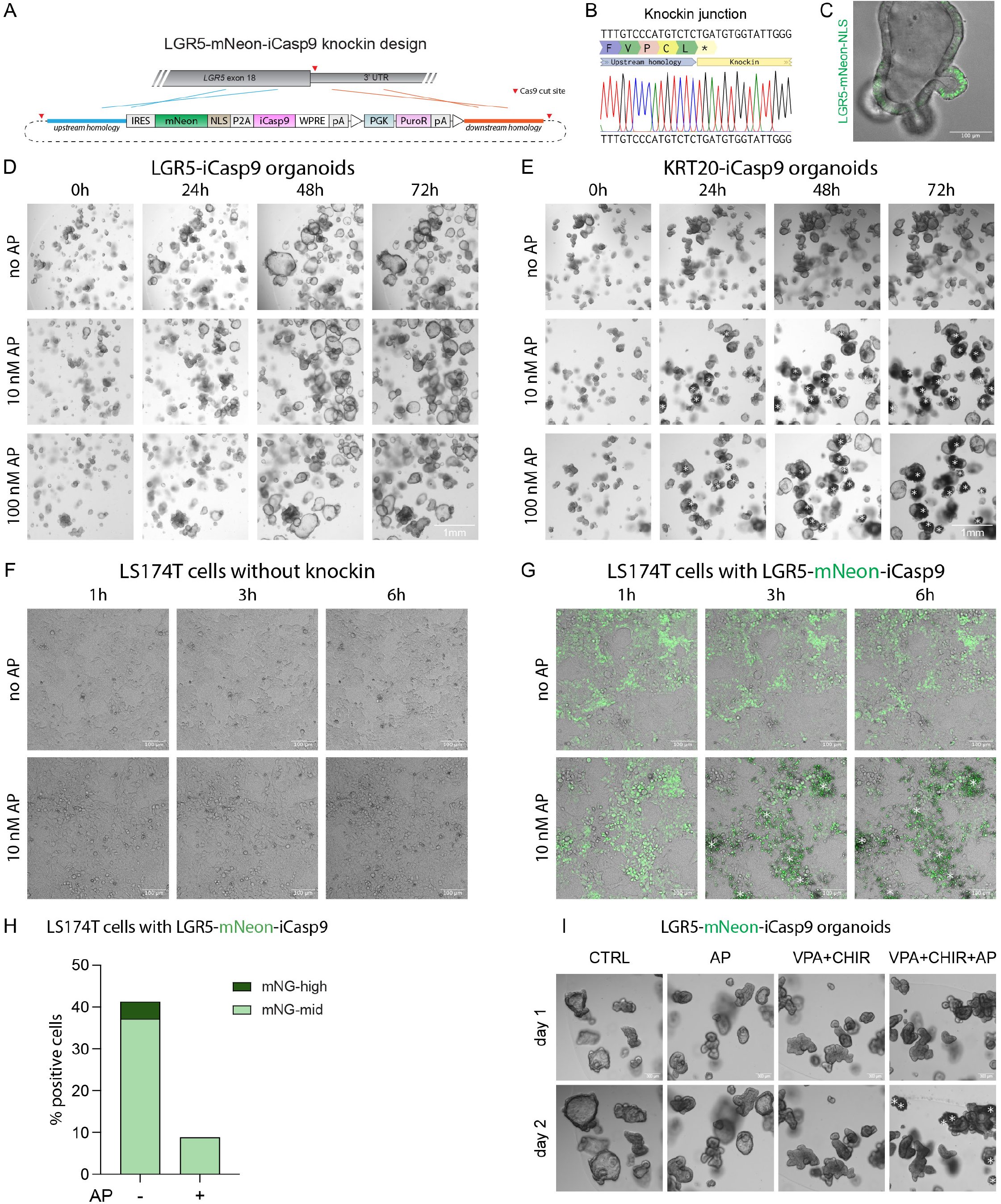
Inducible Caspase 9 is not functional at the *LGR5* locus of normal human colon organoids. **(A)** CRISPR design to generate LGR5-mNeonGreen-inducible Caspase 9 (iCasp9) organoids using in-trans paired Cas9 targeting. Based on Figure S1, an IRES and WPRE were included to achieve the highest reporter expression. IRES: internal ribosome entry site, NLS: nuclear localization signal, WPRE: WHP Posttranscriptional Response Element, pA: poly A sequence, PGK: PGK promoter, PuroR: Puromycin Resistance cassette. **(B)** Sanger sequencing to confirm the genomic knockin integrity of LGR5-mNeon-iCasp9 organoids. **(C)** Microscopy still showing LGR5-mNeonGreen-NLS^+^ cells in the buds of normal human colon organoids. **(D)** Brightfield imaging at indicated timepoints showing that iCasp9 activation with the dimerizer AP20187 (AP) does not cause any observable cell death in LGR5-iCasp9 normal human colon organoids. **(E)** Brightfield imaging at indicated timepoints showing that iCasp9 activation with AP induces cell death in KRT20-iCasp9 normal human colon organoids. Asterisks indicate dying organoids. **(F)** Live cell microscopy showing that LS174T cells without iCasp9 knockin are unperturbed after AP administration. **(G)** Live cell microscopy showing that LGR5-mNeonGreen-iCasp9^+^ LS174T cells are successfully killed upon AP administration. Asterisks indicate areas where cell death is observed. **(H)** Flow cytometry of LS174T cells showing that the majority of LGR5-mNeon-iCasp9^+^ cells is depleted by AP, especially the cells that have high expression levels of mNeonGreen (mNG). **(I)** Brightfield imaging of LGR5-mNeon-iCasp9 normal human colon organoids in different conditions at the indicated timepoints. Asterisks indicate dying organoids.

**Figure S4.**
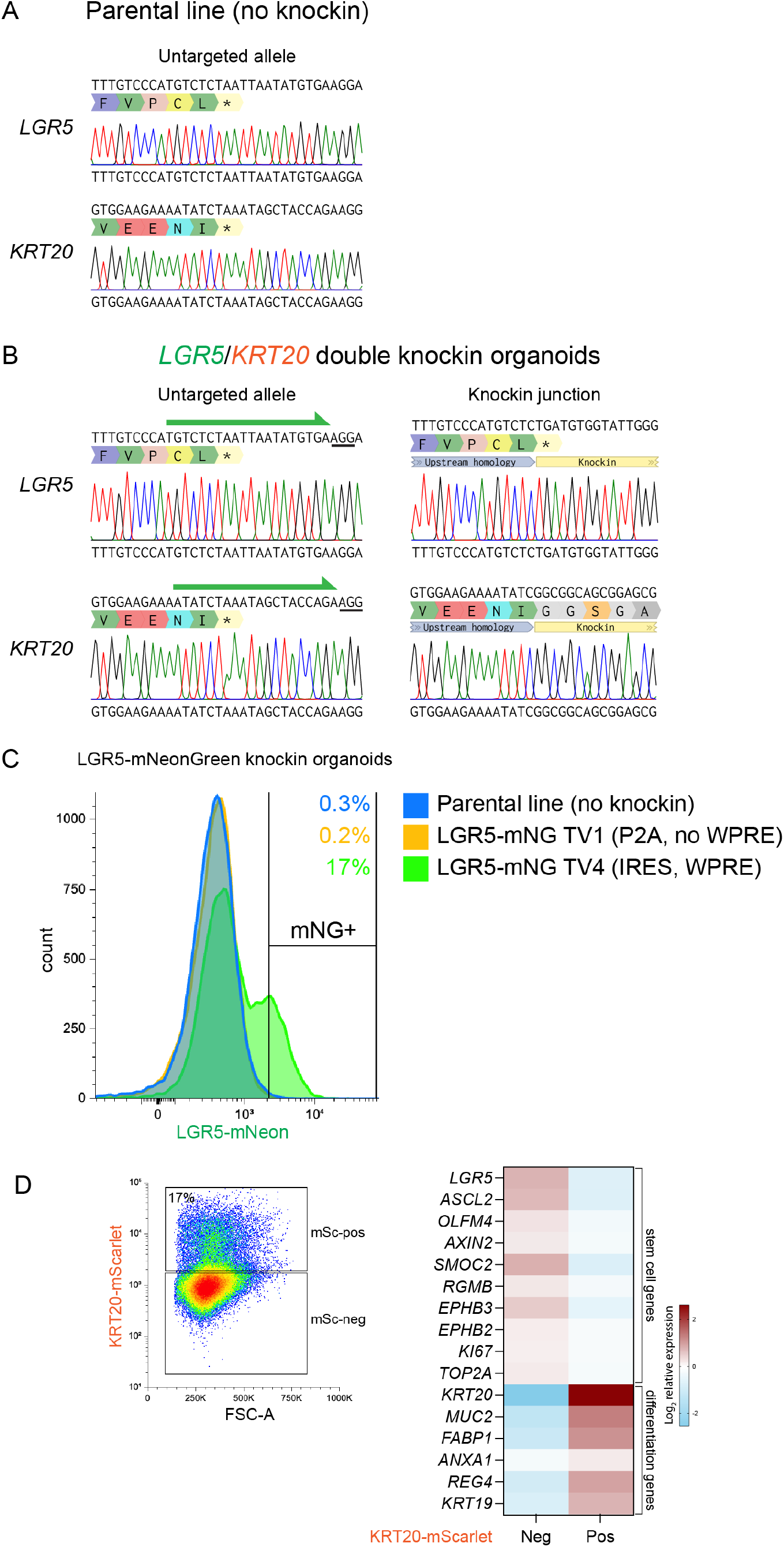
Characterization of LGR5/KRT20 double knockin normal human colon organoids. **(A)** Sanger sequencing of wildtype normal human colon organoids (without knockins). **(B)** Sanger sequencing confirmed the genomic intregrity of LGR5/KRT20 double knockin organoids on the knockin allele as well as the untargeted (secondary) allele. **(C)** Flow cytometry of two CRISPR approaches to knockin mNeonGreen (mNG) at the *LGR5* locus of normal human colon organoids. Targeting Vector 4 (TV4) that includes an IRES and WPRE resulted in the best detectable expression of LGR5-mNG. **(D)** Normal human colon organoids with KRT20-mScarlet (mSc) were FACS sorted (left) and gene expression was analyzed with qPCR (right).

**Figure S5.**
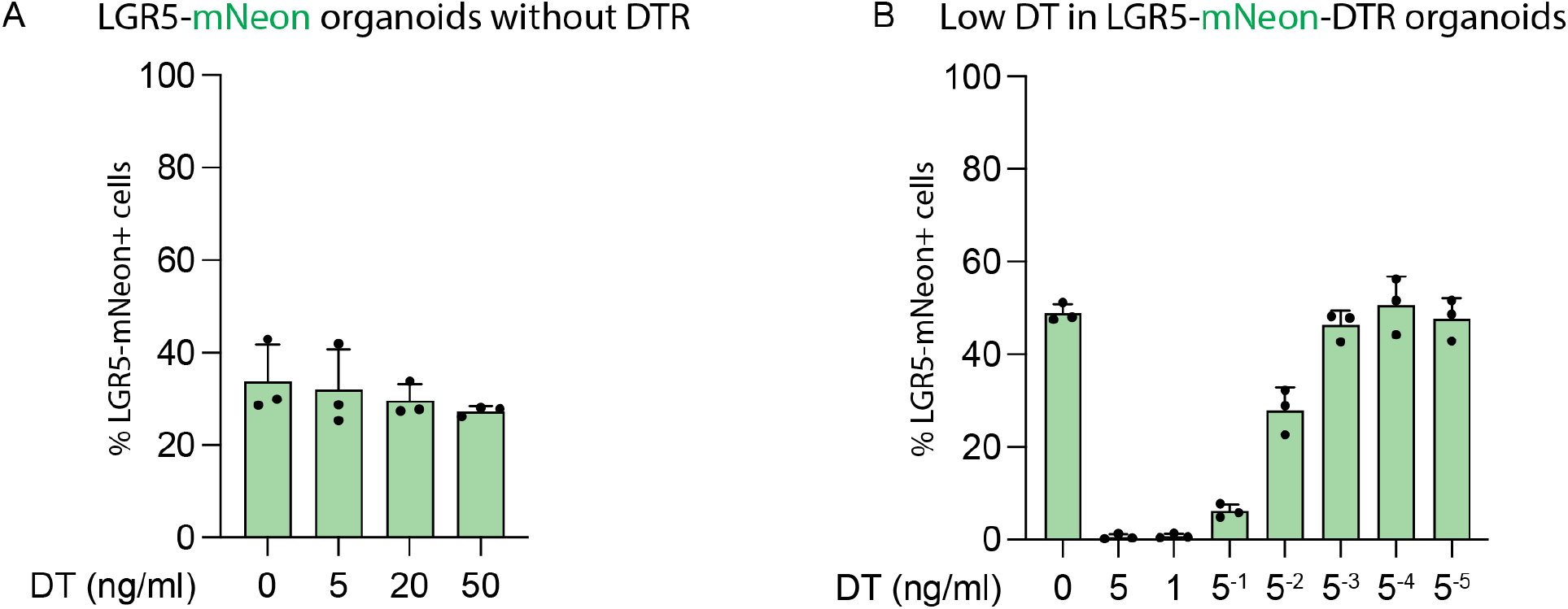
Reappearance of *LGR5*^+^ cells is not an artificial effect of Diphtheria Toxin. **(A)** Flow cytometry of LGR5-mNeon organoids without a DTR, 2 days after a 6-hour DT pulse (when *LGR5*^+^ cell reappearance is normally seen in LGR5-mNeon-DTR organoids). DT does not induce LGR5 expression, even at concentrations 10x higher than used in depletion experiments. Data are mean ± SD. *n* = 3 independent experiments. **(B)** Flow cytometry of LGR5-mNeon-DTR organoids treated for 2 days with low concentrations of DT, showing that low (sublethal) concentrations of DT do not induce expression of *LGR5*. Data are mean ± SD. *n* = 3 independent experiments.

**Figure S6.**
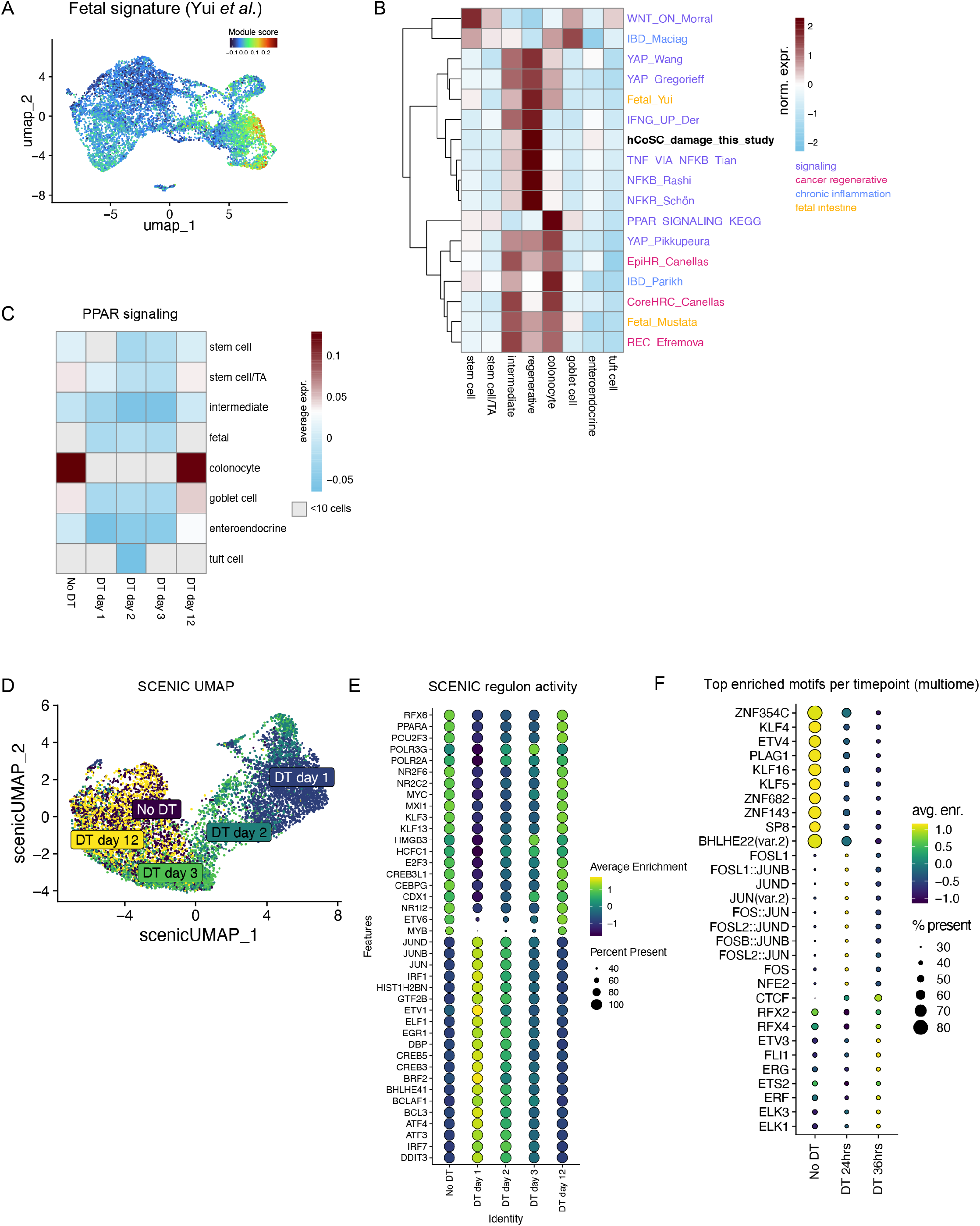
scRNA-seq analysis of the regenerative state following stem cell depletion. **(A)** UMAP visualization of the scRNA-seq dataset before and after stem cell depletion, colored by the activity of the fetal gene signature^8^ **(B)** Heatmap showing the row-normalized expression of the indicated gene signatures per cell type, relating to signaling (purple), regenerative state in cancer (magenta), chronic inflammation (blue), or the fetal intestine (yellow). Signatures were derived from mSigDB, this study, or refs^8,37,51–59^ (see Table S2). **(C)** Heatmap showing the average activity of the PPAR signaling pathway (KEGG) per cell type and timepoint. Groups of cells smaller than 10 were excluded (gray). **(D)** UMAP based on the SCENIC regulon activity scores of the different timepoints. **(E)** Dot plot depicting the top enriched regulons of No DT and DT day 1, displayed for each timepoint. **(F)** Dot plot showing the top enriched motifs per timepoint based on ATAC-seq (ChromVar).

**Figure S7.**
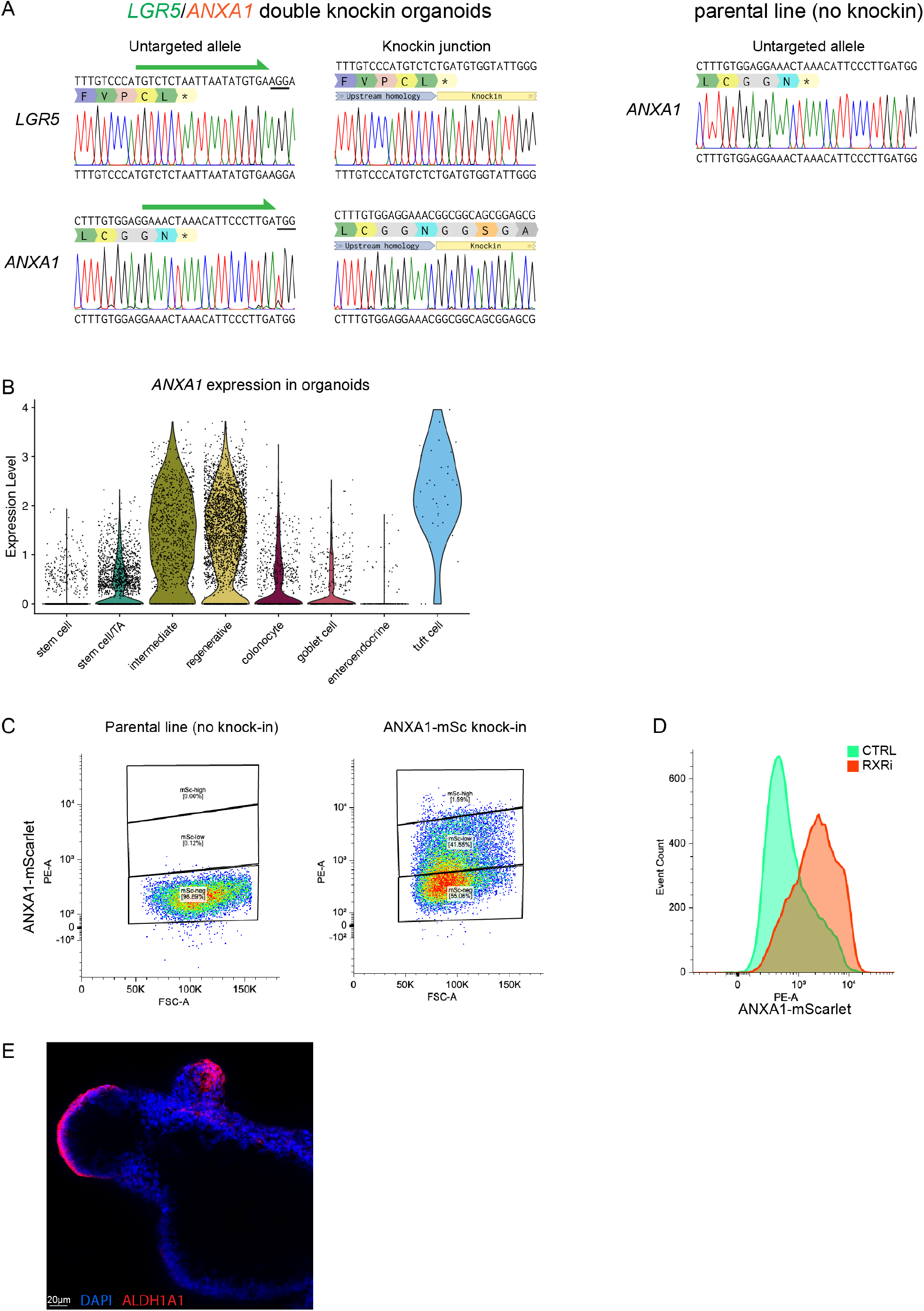
LGR5/ANXA1 double knockin organoids and ALHD1A1 protein expression. **(A)** Sanger sequencing confirming the genomic integrity of LGR5/ANXA1 double knockin organoids on the knockin allele as well as the untargeted (secondary) allele. **(B)** Violin plot depicting the expression of *ANXA1* in normal human colon organoids per cell type in our scRNA-seq dataset, showing high expression in regenerative, intermediate, and (rare) Tuft cells, and low expression in other cell types. Dots represent single cells. **(C)** Flow cytometry gating strategy for ANXA1-mScarlet in normal human colon organoids. **(D)** Flow cytometry histogram relating to Figure 5C showing the shift in *ANXA1* expression upon 24 hrs of RXRi (HX531) treatment. Representative example of *n* = 3 independent experiments. **(E)** Microscopy image of fluorescent antibody staining against ALDH1A1 in normal human colon organoids.

## Notes

### Competing Interest Statement

The authors have declared no competing interest.

## References

1. Barker, N., van Es, J.H., Kuipers, J., Kujala, P., van den Born, M., Cozijnsen, M., Haegebarth, A., Korving, J., Begthel, H., Peters, P.J., et al. (2007). Identification of stem cells in small intestine and colon by marker gene Lgr5. Nature 449, 1003–1007. 10.1038/nature06196.

2. Tian, H., Biehs, B., Warming, S., Leong, K.G., Rangell, L., Klein, O.D., and de Sauvage, F.J. (2011). A reserve stem cell population in small intestine renders Lgr5-positive cells dispensable. Nature 478, 255–259. 10.1038/nature10408.

3. Hageman, J.H., Heinz, M.C., Kretzschmar, K., van der Vaart, J., Clevers, H., and Snippert, H.J.G. (2020). Intestinal Regeneration: Regulation by the Microenvironment. Dev Cell 54, 435–446. 10.1016/J.DEVCEL.2020.07.009.

4. van Es, J.H., Sato, T., van de Wetering, M., Lyubimova, A., Yee Nee, A.N., Gregorieff, A., Sasaki, N., Zeinstra, L., van den Born, M., Korving, J., et al. (2012). Dll1+ secretory progenitor cells revert to stem cells upon crypt damage. Nat Cell Biol 14, 1099–1104. 10.1038/ncb2581.

5. Tetteh, P.W., Basak, O., Farin, H.F., Wiebrands, K., Kretzschmar, K., Begthel, H., van den Born, M., Korving, J., de Sauvage, F., van Es, J.H., et al. (2016). Replacement of lost Lgr5-positive stem cells through plasticity of their enterocyte-lineage daughters. Cell Stem Cell 18, 203–213. 10.1016/j.stem.2016.01.001.

6. Buczacki, S.J.A., Zecchini, H.I., Nicholson, A.M., Russell, R., Vermeulen, L., Kemp, R., and Winton, D.J. (2013). Intestinal label-retaining cells are secretory precursors expressing Lgr5. Nature 495, 65–69. 10.1038/nature11965.

7. Nusse, Y.M., Savage, A.K., Marangoni, P., Rosendahl-Huber, A.K.M., Landman, T.A., De Sauvage, F.J., Locksley, R.M., and Klein, O.D. (2018). Parasitic helminths induce fetal-like reversion in the intestinal stem cell niche. Nature 559, 109–113. 10.1038/s41586-018-0257-1.

8. Yui, S., Azzolin, L., Maimets, M., Pedersen, M.T., Fordham, R.P., Hansen, S.L., Larsen, H.L., Guiu, J., Alves, M.R.P., Rundsten, C.F., et al. (2018). YAP/TAZ-dependent reprogramming of colonic epithelium links ECM remodeling to tissue regeneration. Cell Stem Cell 22, 35–49. 10.1016/j.stem.2017.11.001.

9. Sprangers, J., Zaalberg, I.C., and Maurice, M.M. (2021). Organoid-based modeling of intestinal development, regeneration, and repair. Cell Death Differ 28, 95–107. 10.1038/S41418-020-00665-Z.

10. Serra, D., Mayr, U., Boni, A., Lukonin, I., Rempfler, M., Challet Meylan, L., Stadler, M.B., Strnad, P., Papasaikas, P., Vischi, D., et al. (2019). Self-organization and symmetry breaking in intestinal organoid development. Nature 569, 66–72. 10.1038/s41586-019-1146-y.

11. Sato, T., Stange, D.E., Ferrante, M., Vries, R.G.J., van Es, J.H., van den Brink, S., van Houdt, W.J., Pronk, A., van Gorp, J., Siersema, P.D., et al. (2011). Long-term expansion of epithelial organoids from human colon, adenoma, adenocarcinoma, and Barrett’s epithelium. Gastroenterology 141, 1762–1772. 10.1053/j.gastro.2011.07.050.

12. Ishikawa, K., Sugimoto, S., Oda, M., Fujii, M., Takahashi, S., Ohta, Y., Takano, A., Ishimaru, K., Matano, M., Yoshida, K., et al. (2022). Identification of Quiescent LGR5+ Stem Cells in the Human Colon. Gastroenterology 163, 1391–1406.e24. 10.1053/J.GASTRO.2022.07.081.

13. Fujii, M., Matano, M., Toshimitsu, K., Takano, A., Mikami, Y., Nishikori, S., Sugimoto, S., and Sato, T. (2018). Human intestinal organoids maintain self-renewal capacity and cellular diversity in niche-inspired culture condition. Cell Stem Cell 23, 787–793. 10.1016/j.stem.2018.11.016.

14. Kemper, K., Rodermond, H., Colak, S., Grandela, C., and Medema, J.P. (2012). Targeting colorectal cancer stem cells with inducible caspase-9. Apoptosis 17, 528–537. 10.1007/S10495-011-0692-Z.

15. Shimokawa, M., Ohta, Y., Nishikori, S., Matano, M., Takano, A., Fujii, M., Date, S., Sugimoto, S., Kanai, T., and Sato, T. (2017). Visualization and targeting of LGR5+ human colon cancer stem cells. Nature 545, 187–192. 10.1038/nature22081.

16. Yin, X., Farin, H.F., van Es, J.H., Clevers, H., Langer, R., and Karp, J.M. (2014). Niche-independent high-purity cultures of Lgr5+ intestinal stem cells and their progeny. Nat Methods 11, 106–112. 10.1038/nmeth.2737.

17. Saito, M., Iwawaki, T., Taya, C., Yonekawa, H., Noda, M., Inui, Y., Mekada, E., Kimata, Y., Tsuru, A., and Kohno, K. (2001). Diphtheria toxin receptor-mediated conditional and targeted cell ablation in transgenic mice. Nat Biotechnol 19, 746–750. 10.1038/90795.

18. Naglich, J.G., Metherall, J.E., Russell, D.W., and Eidels, L. (1992). Expression cloning of a diphtheria toxin receptor: identity with a heparin-binding EGF-like growth factor precursor. Cell 69, 1051–1061. 10.1016/0092-8674(92)90623-K.

19. Mitamura, T., Umata, T., Nakano, F., Shishido, Y., Toyoda, T., Itai, A., Kimura, H., and Mekada, E. (1997). Structure-function analysis of the diphtheria toxin receptor toxin binding site by site-directed mutagenesis. J Biol Chem 272, 27084–27090. 10.1074/JBC.272.43.27084.

20. Hooper, K.P., and Eidels, L. (1996). Glutamic acid 141 of the diphtheria toxin receptor (HB-EGF precursor) is critical for toxin binding and toxin sensitivity. Biochem Biophys Res Commun 220, 675–680. 10.1006/BBRC.1996.0463.

21. Aibar, S., González-Blas, C.B., Moerman, T., Huynh-Thu, V.A., Imrichova, H., Hulselmans, G., Rambow, F., Marine, J.C., Geurts, P., Aerts, J., et al. (2017). SCENIC: single-cell regulatory network inference and clustering. Nat Methods 14, 1083–1086. 10.1038/NMETH.4463.

22. Huang, E.H., Hynes, M.J., Zhang, T., Ginestier, C., Dontu, G., Appelman, H., Fields, J.Z., Wicha, M.S., and Boman, B.M. (2009). Aldehyde dehydrogenase 1 is a marker for normal and malignant human colonic stem cells (SC) and tracks SC overpopulation during colon tumorigenesis. Cancer Res 69, 3382–3389. 10.1158/0008-5472.CAN-08-4418.

23. Carpentino, J.E., Hynes, M.J., Appelman, H.D., Tong, Z., Steindler, D.A., Scott, E.W., and Huang, E.H. (2009). Aldehyde dehydrogenase-expressing colon stem cells contribute to tumorigenesis in the transition from colitis to cancer. Cancer Res 69, 8208–8215. 10.1158/0008-5472.CAN-09-1132.

24. Deng, S., Yang, X., Lassus, H., Liang, S., Kaur, S., Ye, Q., Li, C., Wang, L.P., Roby, K.F., Orsulic, S., et al. (2010). Distinct expression levels and patterns of stem cell marker, aldehyde dehydrogenase isoform 1 (ALDH1), in human epithelial cancers. PLoS One 5. 10.1371/JOURNAL.PONE.0010277.

25. Wester, R.A., van Voorthuijsen, L., Neikes, H.K., Dijkstra, J.J., Lamers, L.A., Frölich, S., van der Sande, M., Logie, C., Lindeboom, R.G.H., and Vermeulen, M. (2021). Retinoic acid signaling drives differentiation toward the absorptive lineage in colorectal cancer. iScience 24. 10.1016/J.ISCI.2021.103444.

26. Lukonin, I., Serra, D., Challet Meylan, L., Volkmann, K., Baaten, J., Zhao, R., Meeusen, S., Colman, K., Maurer, F., Stadler, M.B., et al. (2020). Phenotypic landscape of intestinal organoid regeneration. Nature 586, 275–280. 10.1038/S41586-020-2776-9.

27. Vasquez, E.G., Nasreddin, N., Valbuena, G.N., Mulholland, E.J., Belnoue-Davis, H.L., Eggington, H.R., Schenck, R.O., Wouters, V.M., Wirapati, P., Gilroy, K., et al. (2022). Dynamic and adaptive cancer stem cell population admixture in colorectal neoplasia. Cell Stem Cell 29, 1213–1228.e8. 10.1016/J.STEM.2022.07.008.

28. Viragova, S., Li, D., and Klein, O.D. (2024). Activation of fetal-like molecular programs during regeneration in the intestine and beyond. Cell Stem Cell 31, 949–960. 10.1016/J.STEM.2024.05.009.

29. Tierney, M.T., Polak, L., Yang, Y., Abdusselamoglu, M.D., Baek, I., Stewart, K.S., and Fuchs, E. (2024). Vitamin A resolves lineage plasticity to orchestrate stem cell lineage choices. Science (1979) 383. 10.1126/SCIENCE.ADI7342,.

30. Houtekamer, R.M., van der Net, M.C., Maurice, M.M., and Gloerich, M. (2022). Mechanical forces directing intestinal form and function. Curr Biol 32, R791–R805. 10.1016/J.CUB.2022.05.041.

31. Chen, L., Qiu, X., Dupre, A., Pellon-Cardenas, O., Fan, X., Xu, X., Rout, P., Walton, K.D., Burclaff, J., Zhang, R., et al. (2023). TGFB1 induces fetal reprogramming and enhances intestinal regeneration. Cell Stem Cell 30, 1520–1537.e8. 10.1016/J.STEM.2023.09.015.

32. Yan, K.S., Gevaert, O., Zheng, G.X.Y., Anchang, B., Probert, C.S., Larkin, K.A., Davies, P.S., Cheng, Z., Kaddis, J.S., Han, A., et al. (2017). Intestinal enteroendocrine lineage cells possess homeostatic and injury-inducible stem cell activity. Cell Stem Cell 21, 78–90. 10.1016/j.stem.2017.06.014.

33. Ayyaz, A., Kumar, S., Sangiorgi, B., Ghoshal, B., Gosio, J., Ouladan, S., Fink, M., Barutcu, S., Trcka, D., Shen, J., et al. (2019). Single-cell transcriptomes of the regenerating intestine reveal a revival stem cell. Nature 569, 121–125. 10.1038/s41586-019-1154-y.

34. Yan, K.S., Chia, L.A., Li, X., Ootani, A., Su, J., Lee, J.Y., Su, N., Luo, Y., Heilshorn, S.C., Amieva, M.R., et al. (2012). The intestinal stem cell markers Bmi1 and Lgr5 identify two functionally distinct populations. Proc Natl Acad Sci U S A 109, 466–471. 10.1073/PNAS.1118857109.

35. Barriga, F.M., Montagni, E., Mana, M., Mendez-Lago, M., Hernando-Momblona, X., Sevillano, M., Guillaumet-Adkins, A., Rodriguez-Esteban, G., Buczacki, S.J.A., Gut, M., et al. (2017). Mex3a marks a slowly dividing subpopulation of Lgr5+ intestinal stem cells. Cell Stem Cell 20, 801–816. 10.1016/j.stem.2017.02.007.

36. Huang, L., Bernink, J.H., Giladi, A., Krueger, D., van Son, G.J.F., Geurts, M.H., Busslinger, G., Lin, L., Begthel, H., Zandvliet, M., et al. (2024). Tuft cells act as regenerative stem cells in the human intestine. Nature 634, 929–935. 10.1038/S41586-024-07952-6.

37. Cañellas-Socias, A., Cortina, C., Hernando-Momblona, X., Palomo-Ponce, S., Mulholland, E.J., Turon, G., Mateo, L., Conti, S., Roman, O., Sevillano, M., et al. (2022). Metastatic recurrence in colorectal cancer arises from residual EMP1+ cells. Nature 611, 603–613. 10.1038/S41586-022-05402-9.

38. Heinz, M.C., Peters, N.A., Oost, K.C., Lindeboom, R.G.H., van Voorthuijsen, L., Fumagalli, A., van der Net, M.C., de Medeiros, G., Hageman, J.H., Verlaan-Klink, I., et al. (2022). Liver Colonization by Colorectal Cancer Metastases Requires YAP-Controlled Plasticity at the Micrometastatic Stage. Cancer Res 82, 1953–1968. 10.1158/0008-5472.CAN-21-0933.

39. Mzoughi, S., Schwarz, M., Wang, X., Demircioglu, D., Ulukaya, G., Mohammed, K., Zorgati, H., Torre, D., Tomalin, L.E., Di Tullio, F., et al. (2025). Oncofetal reprogramming drives phenotypic plasticity in WNT-dependent colorectal cancer. Nat Genet 57. 10.1038/S41588-024-02058-1.

40. Baas, A.F., Kuipers, J., Van Der Wel, N.N., Batlle, E., Koerten, H.K., Peters, P.J., and Clevers, H.C. (2004). Complete polarization of single intestinal epithelial cells upon activation of LKB1 by STRAD. Cell 116, 457–466. 10.1016/S0092-8674(04)00114-X.

41. Bollen, Y., Hageman, J.H., van Leenen, P., Derks, L.L.M., Ponsioen, B., Buissant des Amorie, J.R., Verlaan-Klink, I., van den Bos, M., Terstappen, L.W.M.M., van Boxtel, R., et al. (2022). Efficient and error-free fluorescent gene tagging in human organoids without double-strand DNA cleavage. PLoS Biol 20. 10.1371/JOURNAL.PBIO.3001527.

42. Buissant des Amorie, J.R., Betjes, M.A., Bernink, J., Hageman, J.H., Heinz, M.C., Jordens, I., Vinck, T., Houtekamer, R.M., Verlaan-Klink, I., Brunner, S.R., et al. (2024). Mature tuft cell phenotypes are sequentially expressed along the intestinal crypt-villus axis following cytokine-induced tuft cell hyperplasia. bioRxiv, 2024.11.28.625899. 10.1101/2024.11.28.625899.

43. Bollen, Y., Post, J., Koo, B.K., and Snippert, H.J.G. (2018). How to create state-of-the-art genetic model systems: strategies for optimal CRISPR-mediated genome editing. Nucleic Acids Res 46, 6435–6454. 10.1093/NAR/GKY571.

44. Ran, F.A., Hsu, P.D., Wright, J., Agarwala, V., Scott, D.A., and Zhang, F. (2013). Genome engineering using the CRISPR-Cas9 system. Nat Protoc 8, 2281–2308. 10.1038/NPROT.2013.143.

45. Fujii, M., Matano, M., Nanki, K., and Sato, T. (2015). Efficient genetic engineering of human intestinal organoids using electroporation. Nat Protoc 10, 1474–1485. 10.1038/NPROT.2015.088.

46. Dekkers, J.F., Alieva, M., Wellens, L.M., Ariese, H.C.R., Jamieson, P.R., Vonk, A.M., Amatngalim, G.D., Hu, H., Oost, K.C., Snippert, H.J.G., et al. (2019). High-resolution 3D imaging of fixed and cleared organoids. Nat Protoc 14, 1756–1771. 10.1038/S41596-019-0160-8.

47. Hao, Y., Stuart, T., Kowalski, M.H., Choudhary, S., Hoffman, P., Hartman, A., Srivastava, A., Molla, G., Madad, S., Fernandez-Granda, C., et al. (2023). Dictionary learning for integrative, multimodal, and massively scalable single-cell analysis. Nat Biotechnol 42, 293. 10.1038/S41587-023-01767-Y.

48. Heaton, H., Talman, A.M., Knights, A., Imaz, M., Gaffney, D.J., Durbin, R., Hemberg, M., and Lawniczak, M.K.N. (2020). Souporcell: robust clustering of single-cell RNA-seq data by genotype without reference genotypes. Nat Methods 17, 615–620. 10.1038/S41592-020-0820-1.

49. Schep, A.N., Wu, B., Buenrostro, J.D., and Greenleaf, W.J. (2017). chromVAR: inferring transcription-factor-associated accessibility from single-cell epigenomic data. Nat Methods 14, 975–978. 10.1038/NMETH.4401.

50. Fornes, O., Castro-Mondragon, J.A., Khan, A., Van Der Lee, R., Zhang, X., Richmond, P.A., Modi, B.P., Correard, S., Gheorghe, M., Baranašić, D., et al. (2020). JASPAR 2020: update of the open-access database of transcription factor binding profiles. Nucleic Acids Res 48, D87–D92. 10.1093/NAR/GKZ1001.

51. Gregorieff, A., Liu, Y., Inanlou, M.R., Khomchuk, Y., and Wrana, J.L. (2015). Yap-dependent reprogramming of Lgr5+ stem cells drives intestinal regeneration and cancer. Nature 526, 715–718. 10.1038/nature15382.

52. Morral, C., Stanisavljevic, J., Hernando-Momblona, X., Mereu, E., Álvarez-Varela, A., Cortina, C., Stork, D., Slebe, F., Turon, G., Whissell, G., et al. (2020). Zonation of Ribosomal DNA Transcription Defines a Stem Cell Hierarchy in Colorectal Cancer. Cell Stem Cell 26, 845–861. e12. 10.1016/J.STEM.2020.04.012.

53. Der, S.D., Zhou, A., Williams, B.R.G., and Silverman, R.H. (1998). Identification of genes differentially regulated by interferon alpha, beta, or gamma using oligonucleotide arrays. Proc Natl Acad Sci U S A 95, 15623–15628. 10.1073/PNAS.95.26.15623.

54. Schön, M., Wienrich, B.G., Kneitz, S., Sennefelder, H., Amschler, K., Vöhringer, V., Weber, O., Stiewe, T., Ziegelbauer, K., and Schön, M.P. (2008). KINK-1, a novel small-molecule inhibitor of IKKbeta, and the susceptibility of melanoma cells to antitumoral treatment. J Natl Cancer Inst 100, 862–875. 10.1093/JNCI/DJN174.

55. Parikh, K., Antanaviciute, A., Fawkner-Corbett, D., Jagielowicz, M., Aulicino, A., Lagerholm, C., Davis, S., Kinchen, J., Chen, H.H., Alham, N.K., et al. (2019). Colonic epithelial cell diversity in health and inflammatory bowel disease. Nature 567, 49–55. 10.1038/S41586-019-0992-Y.

56. Maciag, G., Hansen, S.L., Krizic, K., Kellermann, L., Inventor Zøylner, M.J., Ulyanchenko, S., Maimets, M., Baattrup, A.M., Riis, L.B., Khodosevich, K., et al. (2024). JAK/STAT signaling promotes the emergence of unique cell states in ulcerative colitis. Stem Cell Reports 19, 1172–1188. 10.1016/J.STEMCR.2024.06.006.

57. Mustata, R.C., Vasile, G., Fernandez-Vallone, V., Strollo, S., Lefort, A., Libert, F., Monteyne, D., Pérez-Morga, D., Vassart, G., and Garcia, M.I. (2013). Identification of Lgr5-independent spheroid-generating progenitors of the mouse fetal intestinal epithelium. Cell Rep 5, 421–432. 10.1016/J.CELREP.2013.09.005.

58. Pikkupeura, L.M., Bressan, R.B., Guiu, J., Chen, Y., Maimets, M., Mayer, D., Schweiger, P.J., Hansen, S.L., Maciag, G.J., Larsen, H.L., et al. (2023). Transcriptional and epigenomic profiling identifies YAP signaling as a key regulator of intestinal epithelium maturation. Sci Adv 9. 10.1126/SCIADV.ADF9460.

59. Wang, Y., Xu, X., Maglic, D., Dill, M.T., Mojumdar, K., Ng, P.K.S., Jeong, K.J., Tsang, Y.H., Moreno, D., Bhavana, V.H., et al. (2018). Comprehensive Molecular Characterization of the Hippo Signaling Pathway in Cancer. Cell Rep 25, 1304–1317.e5. 10.1016/j.celrep.2018.10.001.

